# Personalized polygenic risk prediction and assessment with a Mixture-of-Experts framework

**DOI:** 10.1101/2025.09.15.676165

**Authors:** Shadi Zabad, Yue Li, Simon Gravel

## Abstract

With the increasing availability of high quality genomic data from diverse cohorts, polygenic scores (PRS) have become a mainstay of genetic analyses of complex traits and diseases. Despite their proliferation in numerous research domains, a major obstacle to wider adoption in clinical settings has been the well-established heterogeneity in prediction accuracy across a variety of demographic variables, such as age, sex, and genetic ancestry. To address this deficiency, recent research efforts aimed to improve representation in genetic studies and develop stratified PRS inference methods that greatly enhanced accuracy in minority populations. However, with these stratified scores in hand, it remains unclear how to assign the best score, or mixture of scores, for a particular test individual in the clinic. To bridge this gap, we present `MoEPRS`, an ensemble learning method based on the Mixture-of-Experts framework, that blends the stratified scores using personalized mixing weights to predict the target phenotype. In biobank-scale analyses of 7 complex traits in the UK and CARTaGENE biobanks, we show that `MoEPRS` generally provides modest improvements in prediction accuracy over single source PRS models and its predictive performance is maintained across biobanks. Furthermore, we demonstrate practical use cases where the model automatically identifies and adapts to diverse sources of heterogeneity in the data, which allows for evaluating the strengths and weaknesses of current polygenic scores across various cohort strata.

## 1 Introduction

In the past several years, there has been growing interest in incorporating polygenic risk scores (PRS) into various areas of clinical research and practice, including patient risk stratification, detecting gene-by-drug interactions, and designing effective clinical trials [1–3]. However, one of the major roadblocks facing these downstream applications is the documented variability in PRS accuracy across many demographic and clinical variables, including genetic ancestry, age, sex, and socioeconomic status, among others [4–7]. The causes of this poor transferability and their relative importance for each phenotype are still being elucidated, though our evolving understanding point to two major culprits: Allele frequency and Linkage-Disequilibrium (LD) differences across populations and gene-by-environment interactions [8, 9].

The increasing recognition of the PRS transferability problem has spurred a flurry of research across three main axes. First, there has been a considerable push for better and more equitable representation in genetic studies of complex traits and diseases [10–14]. Second, stratified genetic analyses across different cohort variables, such as ancestry or sex, have recently revealed many examples of heterogeneous and group-specific effect sizes or genetic pathways [11, 15–18]. Finally, a wide array of cross-population PRS methods have been developed with the aim of improving variant effect size estimates in each stratum by jointly modeling association data from different ancestry groups [19–24].

While these research efforts have been influential and yielded many important insights, including significant improvements in the accuracy of polygenic scores, they nonetheless do not address several practical concerns. First, given polygenic scores inferred from different strata, biobanks, or cohorts, how do we choose which model, or mixture of models, are best suited for a particular test individual? Second, how can we know if there are other undocumented sources of heterogeneity that might impact the performance of those scores in novel clinical settings?

To address the first concern, recent research focused on ensemble learning techniques that pool together multiple scores to optimize prediction accuracy on a particular validation cohort [19, 25–27]. The technique most popular in the literature, sometimes known as `MultiPRS`, is a form of linear stacking [28], where the predictions of individual PRS models are combined additively with a tunable set of weights [19, 25, 27]. This approach is simple yet powerful, and we expect it to perform competitively on test cases with small, relatively homogeneous cohorts. At the same time, the model does not provide targeted or individual-specific weighing and, in the presence of pronounced variability in the data, it might yield sub-optimal results. As an alternative, a new crop of methods focused on weighing the various scores based on the similarity of the test individual to each training cohort [29–31]. Similarity in this context is often defined in terms of genetic distance or genetic admixture proportions [30, 31]. These approaches, however, are limited in two fundamental ways. Firstly, in some domains, ancestry might not be the dominant source of heterogeneity and it is unclear how to extend these ideas to other cohort variables, such as age or sex, or their combinations. Secondly, there are many factors that can influence PRS accuracy other than the individual’s similarity to the training cohort. This includes considerations such as training sample size, prior choices, and inference method, among others [32–35].

Here, we present a modeling framework that combines the strengths of both approaches for ensemble PRS inference while allowing for the flexibility to identify and adapt to diverse sources of heterogeneity in biobank-scale data. Our approach is based on the Mixture-of-Experts (MoE) ensemble learning technique, which integrates a collection of predictors with a gating mechanism that learns soft partitions of the data, allowing for expert specialization on heterogeneous sub-cohorts [36–38]. In our proposed framework, the gating mechanism is a model that takes as input a set of covariates for each individual, such as their age, sex, and genomic Principal Components (PCs), and outputs a unique set of mixing proportions, which quantify the importance of each PRS for accurately predicting the individual’s phenotype. The whole model is then trained to maximize prediction accuracy on a validation cohort. Thus, the proposed framework supports personalized weighing of polygenic scores while automatically prioritizing scores that improve overall prediction accuracy.

To test this framework on real biobank-scale data, we applied it to analyze 7 complex traits with documented heterogeneity across variables such as age, sex, ancestry, and other latent factors. Our experiments indicate that the Mixture-of-Experts PRS model (`MoEPRS`) provides modest improvement in prediction accuracy over single source PRSs and is competitive with the `MultiPRS` framework, the popular ensemble baseline. Furthermore, our experiments confirm that the gating model learns generally concordant and transferable mappings from covariates to mixture proportions across two independent biobanks [39, 40], a desirable property for the deployment of these models in practical settings. Finally, we demonstrate that the model is readily interpretable and capable of automatically identifying heterogeneities in the data, even in cases where they might not be immediately obvious. To support this latter goal, we present a number of visualization tools and metrics to showcase how the model detects and tackles those sources of variability.

## 2 Material and Methods

### 2.1 Detailed description of the Mixture-of-Experts PRS model

#### 2.1.1 Background and basic model setup

In its classical setup, the Mixture-of-Experts (MoE) model is an ensemble learning framework that combines a collection of predictors with a gating model that weighs their output on a per-sample basis [36–38]. Mathematically, in the context of genetic prediction, the phenotype *y_i_* for individual *i* is modelled as a weighted sum of the predictions of up to *K* experts:

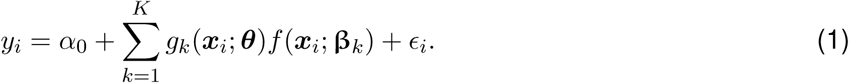

Here, ***x****_i_* is a row vector (1 × *M*) of *M* genetic variants for individual *i*, *α*_0_ is the intercept, and *ɛ_i_* captures the residual effects on the phenotype. *f* (***x****_i_*; **β***_k_*) is the prediction of expert *k* for individual *i* and *g_k_*(***x****_i_*; ***θ***) is its corresponding mixing weight, assigned by the gating model parameterized by ***θ***. Both the mixing weights as well as the predictions are conditional on the individual’s input features ***x****_i_*. What we refer to as mixing weights here can be thought of as probabilities that a phenotype is generated from a given expert, thus satisfying the constraints 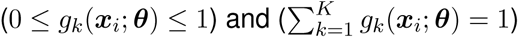 [36, 38]. In this work, we assume that both *g* and *f* are generalized linear functions of the input features:

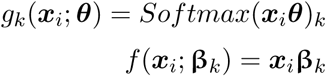

where *θ* is a *M* × *K* matrix of gating model parameters and 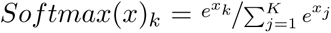 is the Softmax function. For tractability and to scale the model to larger cohorts, we will adopt two major modifications to the classic formulation of the MoE model. First, we let the gating model take as input a different set of features than the experts:

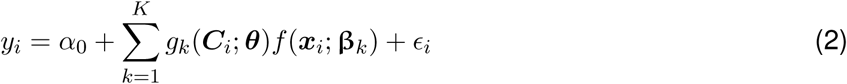

where ***C****_i_* is a row vector of covariates, such as the participant’s age, sex, and top 10 Principal Components (PCs) of the genotype matrix. The main benefit of this modification is that the dimensionality of the covariates is smaller than that of the genotypes. Thus, the gating model will have a substantially smaller set of tuning parameters.

The second modification that we adopt is the assumption that the experts are “pre-trained”, which means that the **β***_k_* are already inferred from diverse, multi-cohort GWAS analyses. For example, the **β***_k_*s could correspond to PRS weights derived from ancestry-, sex-, or age-stratified GWAS and PRS inference results [5, 11, 12]. With these fixed **β***_k_*’s at hand, we can re-write Equation 2 with the following notation:

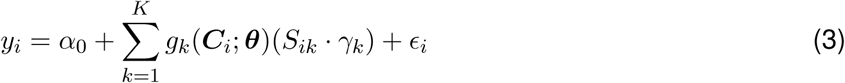

where *S_ik_* := ***x****_i_***β*_k_*** is the polygenic score of sample *i* according to model *k*, and *γ_k_* is a learnable expert-specific scaling factor. In this new formulation, the main task is to train the gating model, parametrized by ***θ***, to optimally combine the pre-trained PRSs for each sample individually.

In early experiments, we found that training the model as outlined in Equation 3 on large multi-ethnic cohorts can lead to poor performance and generalization, mainly due to mean shifts in phenotype and allele frequency differences across populations [41, 42]. To account for these effects when training the `MoEPRS` on diverse cohorts, we augmented the model in Equation 3 to include the covariate effects globally (**Figure** 1) [41, 42],

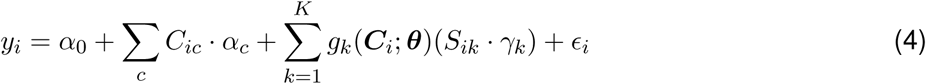

where *C_ic_* is the value of covariate *c* for individual *i* and *α_c_* is the corresponding coefficient. In most of our experiments, these global covariates are the same set of covariates that are fed into the gating model, but in the general case they can differ.

**Figure 1:**
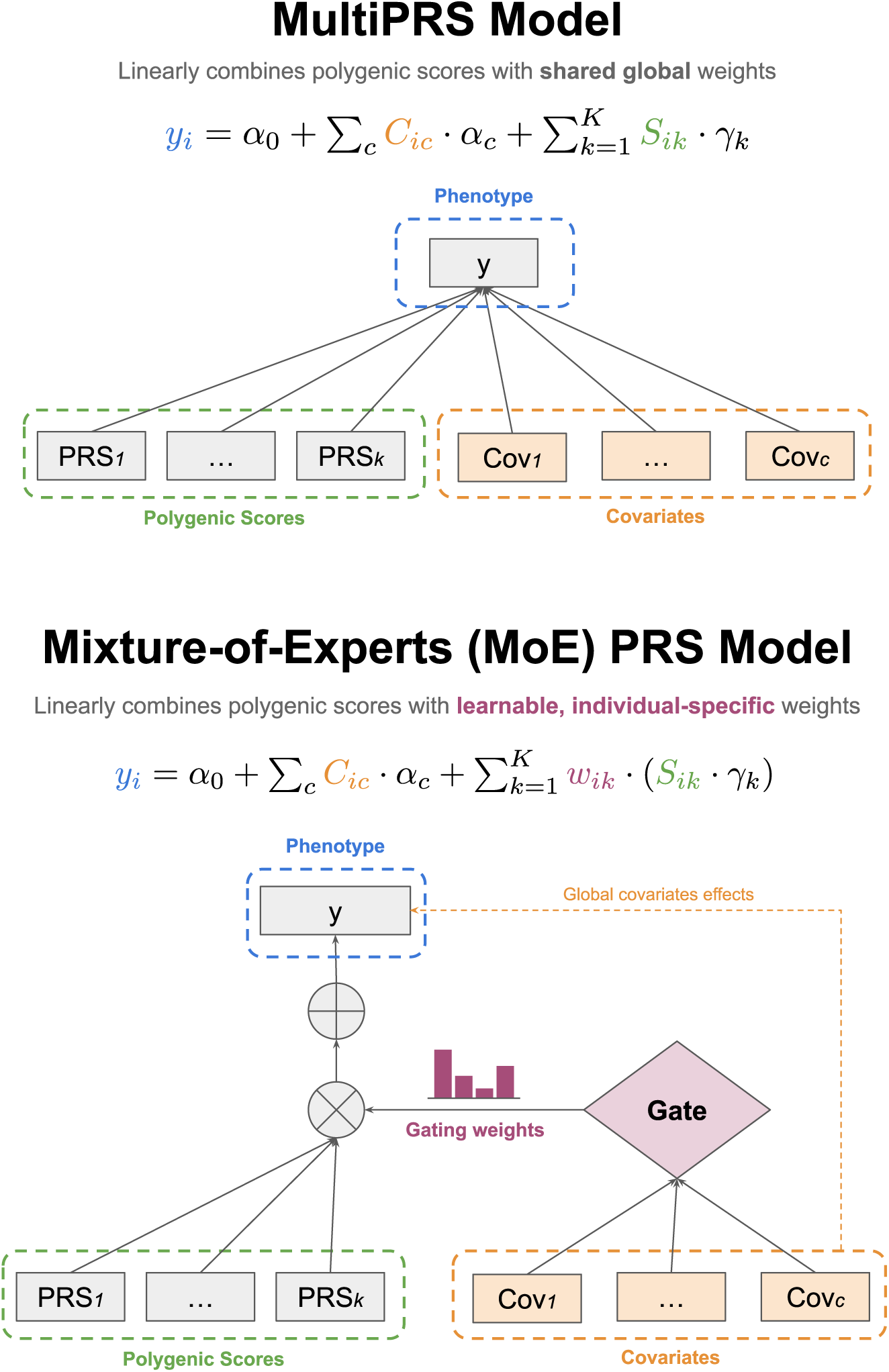
Diagrams illustrating the setup for the ensemble polygenic prediction methods. Standard `MultiPRS` formulations combine single-source polygenic scores linearly, and the weights are shared across the entire cohort. Our proposed `MoEPRS` model combines polygenic scores in a personalized manner, by conditioning the mixing weights on individual attributes, such as age, sex, and genomic Principal Components (PCs).

#### 2.1.2 Fitting Mixture-of-Experts PRS models to biobank data

Assuming we have paired phenotype, genotype, and covariates data from a validation dataset with *N* individuals, training the `MoEPRS` model proceeds according to the following three steps. First, we identify relevant pre-trained polygenic scores for the phenotype of interest, from, e.g. PGS Catalog or similar resources [43, 44]. Second, we use polygenic score weights **β** from the first step to generate the polygenic score matrix, ***S*** = ***X*β**, where ***X*** (*N* × *M*) is the genotype matrix for the validation dataset and **β** (*M* × *K*) is a matrix that contains the PRS weights for *M* variants and *K* different polygenic scores. With the phenotype vector (***y***), PRS matrix (***S***), and the covariates matrix (***C***) at hand, we define the log-likelihood function of the phenotypes as follows:

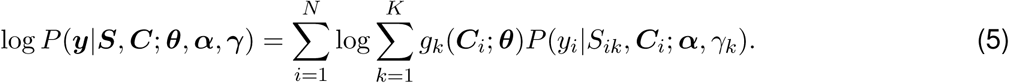

The likelihood function is parametrized by the gating model parameters ***θ***, the shared global covariates parameters *α*, and the expert-specific parameters (e.g. the scaling factors) *γ*. The per-sample likelihood function depends on the category of phenotype considered. For continuous phenotypes, we use Gaussian likelihoods whereas for case-control phenotypes we use Bernoulli likelihood:

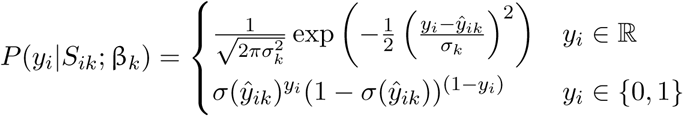

where 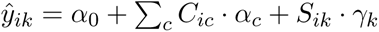 is the scaled and shifted PRS prediction of expert *k* for individual *i* and 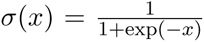 is the logistic function. We implemented two different approaches for fitting the parameters of this model, as outlined below.

##### The EM algorithm

In the classic literature on MoE, the standard procedure for fitting model parameters was the Expectation-Maximization (EM) algorithm [37, 38], an iterative optimization scheme that simplifies the log-likelihood function above by introducing latent indicator random variables that assign each sample to an expert. This “complete-data” log-likelihood then takes the following form:

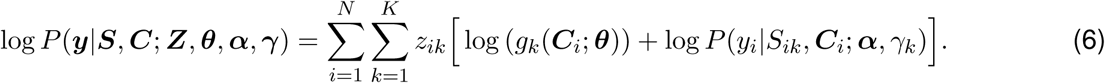

Here, *z_ik_* is a latent categorical variable assigning sample *i* to expert *k*. The main benefit of this approach is that it decouples the gating parameters ***θ*** from the prediction model parameters (***α***, ***γ***), which simplifies the optimization procedure [36, 38]. The EM algorithm proceeds by first initializing the model parameters and then iteratively applying the E- and M-Steps defined below until convergence. For initialization, we randomly set the gating and expert parameters by sampling from a centered Gaussian with scale set to 0.01. The remaining parameters are initialized according to their update equations, outlined below.

In the E-Step, we estimate the expectation for the hidden indicator variables (i.e. “expert responsibility”), given the current values for the gate and expert parameters:

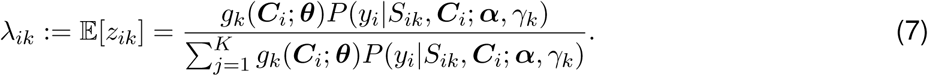

In the M-Step, we optimize the parameters of the gate and the experts conditional on the expert responsibility. To update the gating model parameters, we minimize the cross-entropy between the expert responsibility obtained in the E-Step and the expert weights assigned by the gate,

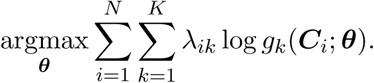

Due to the nonlinearity of the softmax function in *g*(***C****_i_*; ***θ***), there is no closed form solution to this part of the objective function [36, 37]. Therefore, in our implementation, we used the optimizer `L-BFGS-B` from `scipy` [45]. As for the prediction model parameters (***α***, ***γ***), in traditional MoE setups, tuning them is equivalent to fitting a weighted linear model, where the sample weights are specified by the expert responsibilities *λ_ik_* [38]. However, due to the shared parameters ***α***, this is no longer the case and the model requires fitting all the expert weights jointly to maximize the sub-objective:

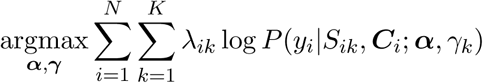

In our implementation, this is again achieved using the L-BFGS-B optimizer from scipy [45].

For continuous phenotypes, the model also infers the residual variance parameters for each expert 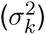 in the M-Step. These parameters are updated using closed-form formulas derived by taking the partial derivatives of Equation 6 [46],

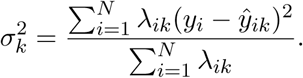

Incorporating expert-specific error variances into the model allows for quantifying the level of uncertainty in their predictions and accommodating heteroscedasticity. In turn, these variance terms affect the expert responsibility, 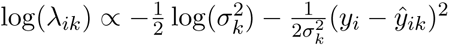. This behavior can be optionally turned off by passing the `fix_residuals` flag to the software, which sets *σ_k_* = 1 for all the experts. In this case, the only factor that affects the partitioning is the squared error of each model, regardless of their prediction uncertainty or differences in phenotypic dispersion patterns.

The EM algorithm described above, along with all the update equations, objectives, and associated derivatives, is implemented in a `python` module that takes the requisite data as input and then performs the iterative EM updates until convergence (**Data and Code Availability**). Convergence is achieved once the difference in the model parameters or the objective function between two consecutive iterations is less than the default tolerance threshold of 10*^−^*^5^. To ensure that the numerical optimization routines are stable, we standardized all continuous variables (covariates, PRSs, and phenotypes) to have zero mean and unit variance before training the model. All the experiments and results in the paper were based on the EM algorithm as outlined above.

##### Stochastic gradient descent (SGD)

The complete data log-likelihood objective discussed above is non-convex and the EM algorithm may be trapped in sub-optimal local minima [37, 38]. Therefore, we also briefly explored an alternative formulation where the model parameters can be jointly optimized to minimize the log-likelihood objective using automatic differentiation libraries, such as `PyTorch` [47]. There are three main benefit to this approach. First, it opens the door for considering more complex gating functions, such as modern neural network architectures. When paired with stochastic optimization techniques, such as the Adam optimizer [48], this can provide two further benefits: It can help scale this algorithm to larger biobank-scale datasets and it may help find better local optima [49]. Our software provides utilities to fit the objectives outlined above using the `PyTorch` library (**Data and Code Availability**), though we did not explore the results from this implementation here.

#### 2.1.3 Prediction in independent test samples

Once the model has been trained on the validation dataset, prediction in independent test samples requires access to the same set of covariates ***C*** used as input for the gating model in addition to the polygenic scores ***S*** for the same set of “experts”. For test sample *i*, their final prediction is given by:

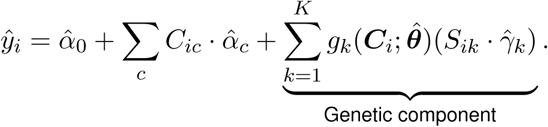

Note that, depending on the phenotype, the linear covariate term can contribute substantially to the prediction. However, in most relevant applications, we are mainly interested in capturing the mixture effect of the genetic component. Therefore, the software provides the option to output predictions based on the mixture of polygenic scores only, omitting the effects of the global covariates.

### 2.2 Baseline methods

To assess the relative improvement provided by the `MoEPRS` model for each phenotype, we compared its predictive performance to a well-established ensemble learning approach known as `MultiPRS` [25, 27]. The `MultiPRS` model was originally proposed as a way to optimize prediction accuracy on underrepresented groups by linearly combining multi-ancestry polygenic scores derived from various cohorts [25]. Mathematically, given a validation dataset with paired genotype, phenotype, and covariates data as above, the `MultiPRS` works by linearly regressing the predictions from multiple polygenic scores against the phenotype (**Figure** 1):

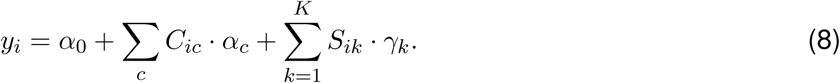

In the case of binary phenotypes, we replace *y_i_* in the equation above with *logit*(*y_i_*). For continuous phenotypes, the `MoEPRS` model reduces to a `MultiPRS` model when the gating model parameters are set to zero (***θ*** = 0). However, without the gating mechanism, the `MultiPRS` is limited since it uses the same global weights for all samples, which may be suboptimal in cases where there are substantial heterogeneities in the data [5], as documented in our results. In principle, one can use nonlinear ensemble methods instead, such as the Boosting machines proposed by Albinana et al. (2022) [26]. However, this can make the model harder to interpret and it may prove more difficult to disentangle the effects of the covariates from those of the genetic predictors.

In our experimental pipeline, we implemented the `MultiPRS` model with the help of `scikit-learn` linear regression modules [50]. Specifically, we provide `python` modules that support linear model fitting with and without regularization penalties, including Ridge, Lasso, and Elastic net. The module takes as input the requisite data (covariates ***C***, PRS matrix ***S***, and phenotype ***y***) and infers the global set of weights ***α*** and ***γ***.

### 2.3 Data processing

#### 2.3.1 Selection of phenotypes and relevant polygenic scores

To examine the performance of the `MoEPRS` model on real phenotype data in modern biobanks, we selected a number of well-studied and significantly heritable phenotypes, with various degrees of documented heterogeneity across several demographic and clinical variables.

The first class of phenotypes included blood biomarker traits with documented heterogeneity by sex [15, 16, 18]: testosterone (UKB Data-field: 30850), creatinine (UKB Data-field: 30700), and urate (UKB Data-field: 30880). To analyze these phenotypes using our framework, we obtained sex-stratified GWAS summary statistics published by Zhu et al. [18] (**Data and Code Availability**) and used the `VIPRS` PRS inference software [51] to infer the polygenic score weights **β** that were used in downstream analyses (**Data and Code Availability**). Note that the GWAS analyses of Zhu et al. primarily used data from “White-British” samples in the UK Biobank [18].

Our second analysis focused on Standing Height (UKB Data-field: 50), a highly heritable and polygenic complex trait, with the largest GWAS sample sizes to date [12]. The comprehensive analyses by Yengo et al. (2022) included ancestry-stratified GWAS for five continental ancestry groups, which was followed by PRS inference using the `SBayesR` software [52]. The ancestry-stratified polygenic scores for the five ancestry groups were obtained from the PGS Catalog (PGP ID: `PGP000382`) [43].

The third and final analysis focused on blood lipid traits, including low-density lipoprotein (LDL) cholesterol (UKB Data-field: 30780), high-density lipoprotein (HDL) cholesterol (UKB Data-field: 30760), and log-transformed triglyceride (UKB Data-field: 30870) levels measured in blood biochemistry assays. These phenotypes are moderately heritable and display various levels of differentiation across age, sex, and ancestry [11, 15, 18]. For LDL cholesterol, we obtained ancestry-stratified polygenic score weights for five continental ancestry groups from the PGS Catalog (PGP ID: `PGP000230`) [11, 43]. For HDL cholesterol and triglycerides, we obtained ancestry-stratified polygenic score weights for four continental ancestry groups from the PGS Catalog (PGP ID: `PGP000489`) [43, 53].

The seven phenotypes and their corresponding polygenic scores, as well as the cohorts from which they were inferred are summarized in **Table** S1. Once the PGS weights were downloaded from the PGS Catalog or other databases, we performed linear scoring on the UKB and CARTaGENE genotype matrices using the `pgsc_calc` (v2.0.0) utility provided by the PGS Catalog researchers [54].

#### 2.3.2 Extraction and pre-processing of UK Biobank data

To apply the `MoEPRS` model to the UK Biobank [39], we extracted genotype, phenotype, and covariates data following standard protocols and Quality Control (QC) procedures. Starting with the genotype data, we extracted records for a total of 486 556 individuals and 2 188 734 variants. We primarily excluded samples who have withdrawn consent, those with inferred high relatedness, and those with putative sex chromosome aneuploidy [39]. For the variants, we kept the union of all the unique variants included in the polygenic scores outlined in the previous section and listed in **Table** S1.

As for the phenotype and covariates data, we extracted raw phenotypic measurements and sample metadata as recorded in the UK Biobank tabular data resource [39]. For each phenotype separately, we excluded samples with outlier values, defined as measurements exceeding 3 standard deviations from the mean. The covariates used in the study include the participant’s age at recruitment, sex, and top 10 Principal Components (PCs). The PCs, along with the polygenic scores ***S***, were computed by passing the `--run_ancestry` flag to the `pgsc_calc` (v2.0.0) software tool [54], which combines the UKB genotype data with that of the harmonized 1000G+HGDP reference samples [55], and projects both jointly onto the same PC coordinates. The `--run_ancestry` flag also fits a random forest classifier that takes the PCs as input and predicts the continental ancestry, as annotated in the 1000G+HGDP resource [54, 55]. The number of samples assigned to each predicted ancestry group are listed in **Table** S2. This predicted ancestry was used in the evaluation and interpretation of the `MoEPRS` model, but not in training. Finally, for an additional layer of interpretability, especially for samples with ambiguous ancestry, we used fine-scale genetic clustering of the UKB samples using the UMAP+HDBSCAN approach outlined recently in Diaz-Papkovich et al. [56].

In our analysis of LDL cholesterol in the UKB, we investigated the relationship between cholesterol-lowering medication and the variability in PGS accuracy by age and sex. To perform this analysis, we extracted self-reported medication records from the tabular data resources from the UKB (Data-field 6177 for males; 6153 for females) [39]. This data was then paired with other demographic variables, such age and sex, to quantify the prevalence of cholesterol drugs across various strata. In follow-up analyses, we also created a new dataset where direct LDL levels were adjusted for medication use. Specifically, if an individual reported taking cholesterol-lowering medication, their measured LDL cholesterol levels were divided by 0.7 as a post-hoc correction [11].

Once all the UKB data sources have been compiled (polygenic scores, phenotypes, covariates, ancestry assignments), we harmonized them using a custom `python` script that combines all of these data into a single dataframe (**Data and Code Availability**). After the data for each phenotype is compiled, we randomly split the samples into 70% training and 30% testing sets.

#### 2.3.3 Extraction and pre-processing of CARTaGENE Biobank data

To examine the robustness and portability of the `MoEPRS` model across biobanks, we extracted genotype, phenotype, and covariates data for the CARTaGENE Biobank (CaG), a prospective cohort from Quebec, Canada [40]. For our downstream analyses, we mainly referenced already pre-processed genotype data used in the CARTaGENE flagship paper [40], which applied all the standard QC criteria for up to 29 333 subjects and included most of the 2 188 734 variants identified earlier.

For the phenotype data, we extracted available phenotypic measurements for 6 of the 7 phenotypes from the tabular data resources. The only phenotype missing from the CaG analysis is testosterone, which is not available in the blood biochemistry results at the time of this publication. For each phenotype, we excluded samples with outlier measurements, which are those exceeding 3 standard deviations from the mean. The covariates used in the study include the participant’s age at recruitment, sex, and top 10 Principal Components (PCs). Similar to what we did in the UKB analysis, the polygenic scores ***S*** and PCs were computed using the `pgsc_calc` (v2.0.0) software utility by combining the CaG genotypes with the 1000G+HGDP reference set and projecting both onto the same PC coordinates [54, 55]. Similarly, this pipeline also generated predicted continental ancestry assignments, which are listed in **Table** S2 and used in all our downstream analyses.

Once all the CaG data sources have been compiled (polygenic scores, phenotypes, covariates, ancestry assignments), we harmonized them using a custom `python` script that combines all of this data into a single dataframe (**Data and Code Availability**). After the data for each phenotype is compiled, we randomly split the samples into 70% training and 30% testing sets.

### 2.4 Model evaluation and interpretation

#### 2.4.1 Metrics for evaluating prediction accuracy

After the models were fit to the training data in each biobank separately, they were assessed on the held-out test sets using standard evaluation metrics. For continuous phenotypes, the main evaluation metric used in the analysis is the Incremental R-Squared, defined as the difference in the proportion of explained variance between a fully parameterized model (*y* ∼ *PRS* + *covariates*) and a null model (*y* ∼ *covariates*). The *PRS* variable here is the prediction, obtained from either single source or ensemble PRS models. To obtain confidence intervals for the R-Squared evaluation metrics, we implemented the recently proposed estimators for the sampling variance of the R-Squared metric [57]. All of these evaluation metrics and quantities can be computed using the latest version of our python package `viprs` (v0.1) [51, 58].

Given that our `MoEPRS` model is not directly optimizing for the incremental *R*^2^ metric, for sanity checking we also used the Pearson Correlation Coefficient and the Mean Squared Error (MSE) between the PRS and the phenotype to assess the quality of the predictions on certain sub-cohorts in the dataset.

#### 2.4.2 PGS admixture graphs

To aid in interpreting the output of the gating mechanism in the `MoEPRS` model, we propose using a modified version of the classic admixture bar plots popularized by the STRUCTURE/ADMIXTURE ancestry inference methods [59–61]. In our application, the mixing proportions for each individual do not represent ancestry assignment, but mixing weights assigned by the gating model to each PGS, conditional on the input covariates ***C****_i_* and the learned parameters 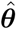 for a particular phenotype. To generate a PGS admixture graph for independent test samples, we first assemble their covariates into a matrix ***C*** (*N* × *C*), where each row contains the gating model covariates for single test sample. In our experiments, these covariates mainly consist of the participant’s age at recruitment, sex, and position along the first 10 Principal Components. The matrix of covariates is then passed to the gating model, which outputs a (*N* × *K*) matrix of mixing weights (i.e. ***Q*** matrix [59, 60]) that are specific to the phenotype in question. *K* here depends on the number of polygenic scores used in the analysis. By construction, the weights are proportions, meaning that their values are between 0 and 1 and each row sums to 1 exactly.

To enhance interpretability of the PGS admixture plots, we categorized the samples by their predicted continental ancestry [55] or UMAP+HDBSCAN cluster membership [56]. Within each group or cluster, the samples are sorted by their mixing proportions for improved visibility.

#### 2.4.3 Concordance of learned PGS admixture proportions

To quantify the agreement between the PGS admixture proportions produced by different `MoEPRS` models for the same phenotype, such as those trained on two distinct biobanks, we first generate the matrix of mixing proportions ***Q*** on the same held-out test set. Once matrices of mixing proportions are generated from each model (e.g. ***Q***_UKB_ and ***Q***_CaG_ for UKB- and CaG-trained models), we can assess their calibration and concordance in a number of ways. Principally, in our analyses, we quantified their concordance by looking at the Pearson Correlation Coefficient between the columns of those matrices, e.g. *R* (***Q***_UKB_(*i*), ***Q***_CaG_(*i*)), where ***Q***(*i*) denotes taking the *i*−th column of ***Q***. For PGSs that have high explanatory value, and consequently, receive substantial weights by the gating model, we expect a strong, significant positive correlation. On the other hand, if the PGS has minor importance and its main use (if any) is to correct for the mistakes of the others, then we expect the correlation to be negligible.

## 3 Results

### 3.1 MoEPRS accommodates diverse sources of heterogeneity in sex-differentiated phenotypes

To illustrate the capabilities of the Mixture-of-Experts framework, we start with a relatively simple setup, where the ensemble PRS model is tasked with combining predictions from two base PGS models derived from male and female participants separately (“Male PGS” and “Female PGS”). Specifically, we focus on moderately heritable blood biomarker traits that have been shown to display various levels of differentiation across the sexes [15, 16, 18]: testosterone, creatinine, and urate. After obtaining sex-specific PGSs for the UK Biobank participants [39] for each of these phenotypes (**Methods**), we trained the `MoEPRS` model to combine them in a personalized manner, where the gating mechanism is conditioned on a set of covariates: The participant’s sex, age at recruitment, and 10 genomic Principal Components (**Figure** 1). For comparison, we also include the `MultiPRS` model [25, 27], which combines the scores using a global set of weights (**Figure** 1). The ensemble PRS models were fit on a training subset comprising 70% of samples in the UKB, irrespective of ancestry or sex.

On a highly sex-differentiated trait like testosterone, the sex-specific PGS models perform well within sex, but generalize poorly to the opposite sex (**Figure** 2A). This is to be expected, as the heritability and genetic architecture of testosterone are highly divergent across the sexes (*r_g_* ≈ 0.1) [15, 16, 18, 62]. Notably, males exhibit much greater variability in testosterone levels than females [16], which explains why, in this setup, the `MultiPRS` model assigns most of its weight to the “Male PGS” and inherits its error profile (**Figure** 2A). The `MoEPRS` model, on the other hand, detects that the error profiles of the two models are correlated with the participant’s sex and uses the gating mechanism to partition the cohort and assign each individual to their sex-matched PGS (**Figure** 2D; S1). This partitioning is uniform across ancestries and age groups (**Figure** 2D;S3A), confirming that gene-by-sex interactions are indeed the dominant factor in explaining the heterogeneity of testosterone levels [15, 16, 18, 62].

**Figure 2:**
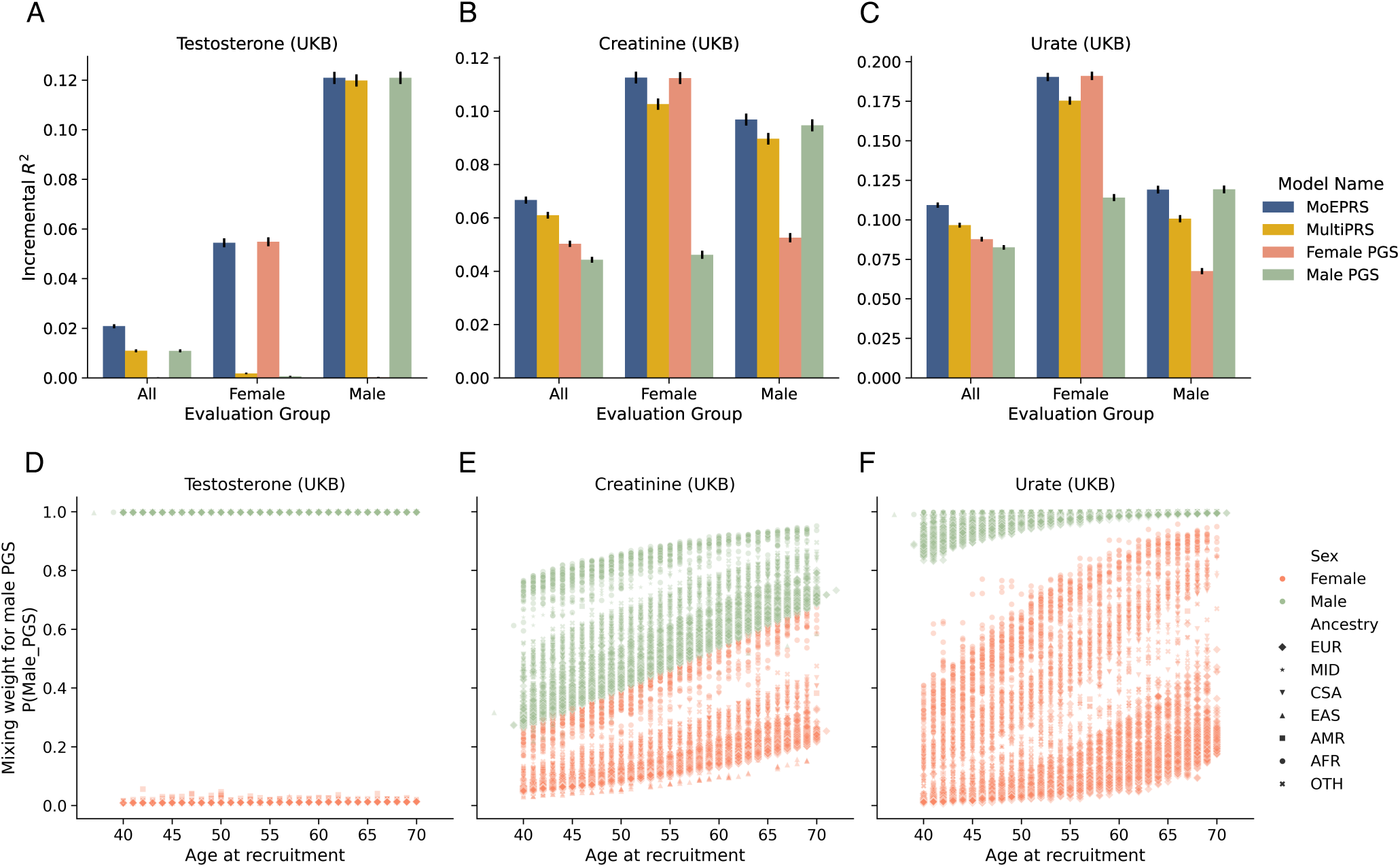
Prediction accuracy and specialization behavior of `MoEPRS` on sex-differentiated complex traits in the UK Biobank. Sex-stratified polygenic scores (“Male PGS” and “Female PGS”) derived from European samples were used as input to predict phenotypes for the entire UKB cohort. Ensemble PRS models that combine the stratified scores include our proposed `MoEPRS` as well as `MultiPRS`, which learns a linear combination of individual scores. The complex traits analyzed are testosterone, creatinine, and urate levels measured in blood biochemistry assays. Panels (**A-C**) show prediction accuracy in terms of incremental *R*^2^ on the held-out test set in the UKB. The accuracy measures are stratified by sex (x-axis), and vertical black lines on top of the bars show analytical standard errors for the *R*^2^ metric. Panels (**D-F**) show the weight assigned to the “Male PGS” by the gating model for each individual in the test set, as a function of the sample’s age at recruitment (x-axis). The scatter points are differentiated by colors (green for males and orange for females) and markers corresponding to different continental ancestry groups. The ancestry groups include: `EUR` (European), `MID` (Middle Eastern), `CSA` (Central and South Asian), `EAS` (East Asian), `AMR` (Admixed American), `AFR` (African), and `OTH` (Other).

In contrast to testosterone, creatinine and urate levels in the blood exhibit stronger genetic correlation across the sexes (*r_g_* ≈ 0.9), in addition to being significantly more heritable in females [15, 18]. As a result, while the predictive performance of the sex-specific PGS models drops when evaluated on the opposite sex, the differences are not as drastic (**Figure** 2A-C). This relative concordance between the sex-specific PGSs allows the `MultiPRS` model to pool together both predictors effectively and improve its performance on the entire sample, though it still falls short on each cohort individually (**Figure** 2B-C). The `MoEPRS` model, on the other hand, detects that sex is again a major driver of heterogeneity for these traits, though remarkably, it identifies other variables that affect the accuracy of the sex-specific models (**Figure** 2B-C). Primarily, their predictive performance seems to vary with age and ancestry (**Figure** S5): the gating model assigns relatively higher weight to the “Male PGS” for older individuals and for certain minority ancestry groups (**Figure** 2E and F; **Figure** S3). This heterogeneity by age and ancestry replicated in an independent analysis of equivalent data in the CARTaGENE biobank (**Figure** S2, S5), a prospective cohort from Quebec, Canada [40].

The age effect observed for creatinine may be explained by noting that, while generally higher in males, average creatinine levels and their dispersion rise with age in both sexes, affected by factors such as muscle mass loss, kidney function, and hormonal changes [63, 64] (**Figure** S4). We confirmed that the “Female PGS” indeed generalizes better to younger individuals across sexes and ancestries, though this advantage generally reverses or vanishes in older samples (**Figure** S5A). As for the observed heterogeneity by ancestry, we hypothesize that they are partly driven by differences in baseline creatinine levels, with samples of African and South Asian ancestry having the highest mean and variance [64] (**Figure** S5A). For urate, the age effect might similarly be attributed to the fact that, while males generally have higher serum urate levels than females, the gap narrows somewhat with age, particularly after menopause [65, 66]. As for the observed cross-ancestry effect, previous studies have documented that individuals of African and Asian ancestry have higher baseline levels of urate [67], and our UKB analysis showed that older females of African and South Asian ancestry exhibit the highest variability in serum urate levels (**Figure** S5B). We also confirmed that the “Male PGS” for urate does transfer better to older females from some ancestry groups in both biobanks (**Figure** S5).

### 3.2 Consistency and portability of MoEPRS on standing height across two biobanks

The next test case for the `MoEPRS` framework is predicting standing height, a highly heritable and polygenic trait with many well-powered, ancestry-stratified polygenic scores [12]. In the largest GWAS study of height to date, Yengo et al. (2022) [12] found that variant effect sizes estimated in five continental ancestry groups are highly correlated (*r_g_* ∈ [0.64, 0.99]). Yet, the corresponding polygenic scores did not generally transfer well across ancestries [12]. To examine how the `MoEPRS` handles those scores in practice, we separately fit the model to height data from the UK and CARTaGENE biobanks [39, 40]. In each case, the gating model took as input the same set of covariates as in the previous experiments: The participant’s age, sex, and 10 PCs. The model was independently trained on a 70% randomly sampled subset in each biobank, irrespective of ancestry.

We examined the prediction accuracy on the held-out test set in each biobank, stratified by inferred continental ancestry group (**Figure** 3). We make three main observations about these results. First, while the ancestry-matched score generally corresponds to the best single source score, it significantly underperforms for Central and South Asian (`CSA`) Admixed American (`AMR`) samples (**Figure** 3A, B). This is consistent with the results of Yengo et al. (2022) and may be partly explained by noting that the these PGSs had the smallest training sample sizes [12]. This finding highlights a limitation of ensemble methods that assign PGSs purely based on the individual’s genetic similarity with the training cohort [30,31] (**Discussion**). Second, in this experiment, `MoEPRS` does better than `MultiPRS` or any single source PGS in aggregate. On the other hand, within subpopulations, the model is on par with the best PGSs in most populations, but underperforms in some cases, e.g. `MoEPRS` on `AMR` samples in UKB (**Figure** 3A). We followed up on this particular case and found that the `MoEPRS` model mostly assigns samples with Admixed American (`AMR`) ancestry to the `EAS` PGS (**Figure** 3D), which indeed has the best performance in terms of marginal Pearson correlation, but once we correct for the effect of the covariates, this model’s relative performance drops precipitously (**Table** S3; see **Discussion**). Finally, despite differences in the ancestry composition of their training data, the `MoEPRS` models generally transfer well within and across biobanks (**Figure** 3A-B), with some minor exceptions.

**Figure 3:**
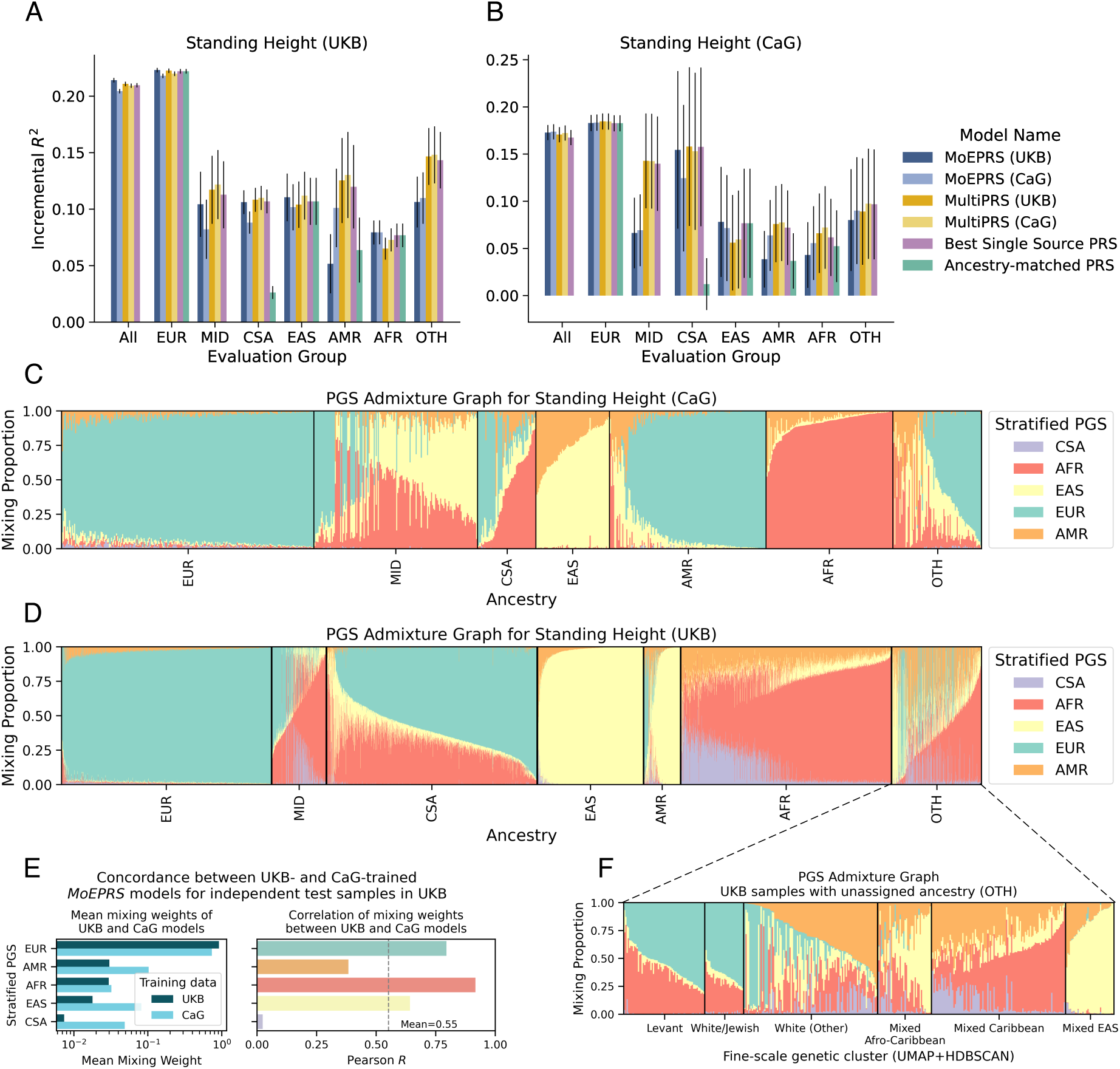
Prediction accuracy and specialization behavior of the `MoEPRS` model on Standing Height in the UK and CARTaGENE biobanks. Ancestry-stratified polygenic scores derived from five continental ancestry groups were used as input to predict height in each biobank separately. Ensemble PRS models that combine the stratified scores include our proposed `MoEPRS` as well as the popular baseline `MultiPRS`, which learns a linear combination of individual scores. Panels (**A-B**) show prediction accuracy in terms of incremental *R*^2^ on the held-out test set in each biobank. The accuracy measures are stratified by ancestry (x-axis), and vertical black lines on top of the bars show analytical standard errors for the *R*^2^ metric. Parentheses in ensemble PRS model name denote the training cohort (UKB or CaG). Panels (**C-D,F**) show PGS admixture graphs, which represent the mixing weights assigned by the gating model for a random subsample of individuals. Panel (**C,D**) show the mixing proportions for independent test samples in the UK and CARTaGENE biobanks, respectively, stratified by continental ancestry. Panel (**F**) shows the mixing proportions for UKB samples with unassigned ancestry (Ancestry = `OTH`), stratified by fine-scale genetic clusters. Panel (**E**) shows the mean mixing weights (left sub-panel) and correlation of mixing weights (right sub-panel) for the `MoEPRS` models trained on the two biobanks and applied to the test samples in the UKB.

The portability of the `MoEPRS` model across biobanks is driven by the consistent mixture patterns learned by the gating models in each biobank (**Figure** 3C, D; **Figure** S6). From the error patterns of each PRS, the gating mechanism automatically learned to mostly assign ancestry-matched PRSs for Europeans (`EUR`), East Asians (`EAS`), and Africans (`AFR`), and settled on similar admixture profiles for the other minority ancestry groups (**Figure** 3C, D). Quantitatively, we see strong correlation in the PGS mixing weights learned by the two models (mean Pearson *R* = 0.55), especially for the three PRSs with the most dominant weights (**Figure** 3E; **Figure** S8). One of the main advantages of the `MoEPRS` for analyzing large-scale and diverse biobanks is that it is not restricted by declared or inferred ancestry and it can learn consistent patterns for individuals with ambiguous ancestry assignments (**Figure** 3F). Furthermore, the model sometimes identifies consistent variation within ancestry labels, for instance detecting different optimal PGS assignment between Levantine and North African samples (**Figure** S7), who are commonly lumped under the “Middle Eastern” (`MID`) group by popular ancestry predictors [55].

### 3.3 MoEPRS identifies trait-specific admixture profiles for blood lipid phenotypes

Lastly, we conducted a detailed analysis of three blood lipid phenotypes whose genetic underpinnings have been systematically investigated in large scale, multi-ancestry cohorts [11]: Low- and High-Density Lipoprotein (HDL and LDL) as well as log-transformed Triglyceride levels. Recent analyses from the Global Lipids Genetics Consortium (GLGC) have shown that these phenotypes are moderately heritable, somewhat polygenic, and influenced by many large effect variants, some of which are ancestry-specific [11]. In general, ancestry-stratified GWAS analyses revealed that variant effect sizes are highly correlated for all three phenotypes and across most ancestry groups, though there are some notable variations (*r_g_* ∈ [0.473, 1]) [11]. To see how the `MoEPRS` model tackles the unique variations underlying each phenotype, we obtained ancestry-stratified polygenic scores and trained the model on harmonized data in the UK and CARTaGENE biobanks [39, 40]. For every phenotype and biobank separately, the model was trained on data for a random of 70% of the samples the gating model received as input the participant’s age, sex, and 10 genomic PCs.

We first assessed the prediction accuracy for all three phenotypes, evaluated on the held out test set in the UK Biobank (**Figure** 4A-C). Unlike standing height, for the blood lipid phenotypes we do see that the ensemble PRS models provide modest improvement in prediction accuracy over the single source PRS, especially on samples of European ancestry. By contrast with standing height, the gating mechanism here distributes the weight across multiple models, instead of favoring a single PRS (**Figure** 4D-F). This mixing pattern is qualitatively reproducible across both biobanks, with some slight differences for some minority groups (**Figure** 4D-F; **Figure** S10D-F). Despite this qualitative similarity, we do see significant drop in the portability of CaG-trained `MoEPRS` model (**Figure** 4A). We presume that this is the result of overfitting on the smaller CARTaGENE dataset, since the transferability of the UKB-trained model is unaffected (**Figure** S10A-C; see **Discussion**). To understand better the blending patterns observed in the PGS admixture graphs (**Figure** 4D-F) and the underlying variables driving them, we investigated the dominant trends for each phenotype separately.

**Figure 4:**
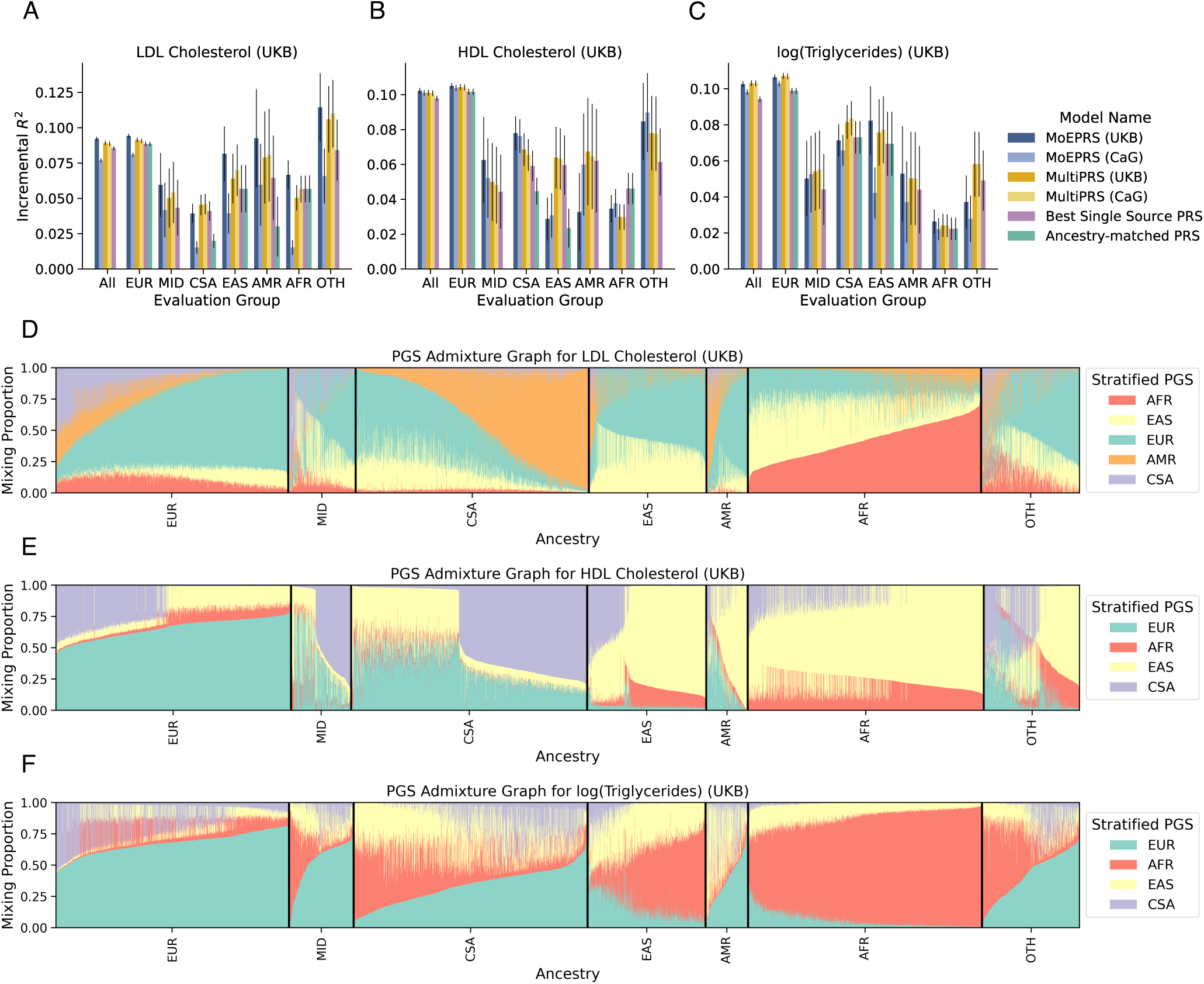
Prediction accuracy and specialization behavior of the `MoEPRS` model on blood lipid traits in the UK Biobank (UKB). Trait-specific and ancestry-stratified polygenic scores derived from up to five continental ancestry groups were used as input to predict each molecular phenotype. The traits analyzed are low-density lipoprotein (LDL) cholesterol, high-density lipoprotein (HDL) cholesterol, and log-transformed triglyceride levels in the blood. Ensemble PRS models that combine the stratified scores include our proposed `MoEPRS` as well as the popular baseline `MultiPRS`, which learns a linear combination of individual scores. Panels (**A-C**) show prediction accuracy in terms of incremental *R*^2^ on the held-out test set in the UKB. The accuracy measures are stratified by ancestry (x-axis), and vertical black lines on top of the bars show analytical standard errors for the *R*^2^ metric. Panels (**D-F**) show PGS admixture graphs for each phenotype, which represent the mixing weights assigned by the gating model for a random subsample of individuals, stratified by continental ancestry.

In the analysis of LDL cholesterol, we found that the main factors driving the variation in PGS admixture proportions in Europeans are the participant’s age and sex (**Figure** 5A; **Figure** S9). Notably, our results show that the `EUR` PGS for LDL performs better in females and its accuracy decreases substantially with age (**Figure** 5B). This variability in prediction accuracy with age and sex was replicated in our analysis of the CARTaGENE biobank (**Figure** S11C). Previous work has shown that LDL cholesterol is significantly more heritable in females [15], which might explain some of the heterogeneity across sex. The evidence for declining heritability with age, on the other hand, is either inconclusive or the effect is too weak to account for this disparity [68, 69]. We hypothesize that the main underlying factor driving this variability is the prevalence of cholesterol lowering medication, which we did not account for in our original analysis. Indeed, age and sex groups with higher prevalence of cholesterol-lowering medication attain lower prediction accuracy (**Figure** 5B). In the original GLGC study, the authors corrected for medication use by dividing the direct LDL cholesterol levels measured in those samples by 0.7 [11]. In our case, since we did not apply this correction, the model identified features correlated with medication use (age and sex) and attempted to improve its accuracy on those samples by blending in information from other polygenic scores. To verify that medication use is indeed the primary factor in explaining this heterogeneity, we repeated the analysis with the adjusted LDL cholesterol phenotype and found that the mixing weights were substantially less variable with age and sex (**Figure** S14;**Figure** S15).

**Figure 5:**
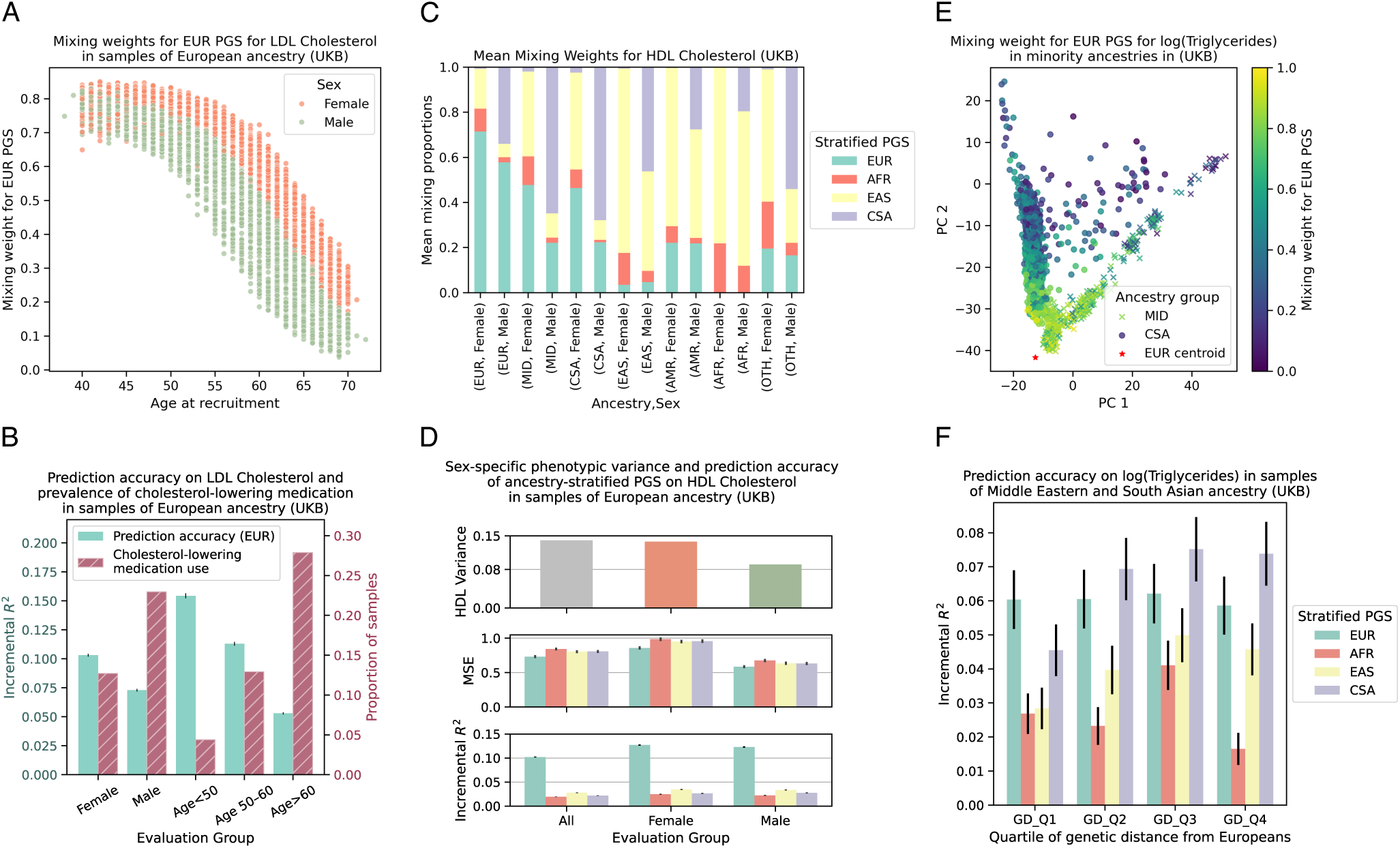
Investigation of a subset of the soft cohort partitions identified by `MoEPRS` in the analysis of blood lipid phenotypes in the UK Biobank. Panel (**A**) shows the mixing weight assigned to the European-derived PGS (`EUR`) model for LDL cholesterol as a function of the sample’s age at recruitment (x-axis) and sex (color). Panel (**B**) shows the prediction accuracy of the `EUR` PGS for LDL Cholesterol (left y-axis) and prevalence of cholesterol-lowering medication (right y-axis) in European samples in the UKB, stratified by sex and age groups (x-axis). Panel (**C**) shows the average mixing proportions assigned by `MoEPRS` to each group of individuals when predicting HDL cholesterol, categorized by the sample’s predicted continental ancestry and sex (x-axis). Panel (**D**) shows, for each sex, the phenotypic variance of HDL cholesterol as well as the predictive performance of the ancestry-stratified PGSs on European samples in the UKB. The predictive performance metrics include Mean Squared Error (MSE) of the covariates-augmented models as well as Incremental *R*^2^. Panel (**E**) shows the mixing weight assigned to the European-derived PGS (`EUR`) for log(Triglycerides) in samples of Middle Eastern and South Asian ancestry, as a function of the individual’s location along the first two genomic PCs. Panel (**F**) shows the prediction accuracy, in terms of incremental *R*^2^, of the various PGSs in Middle Eastern and South Asian samples in the UKB, stratified by quartiles of genetic distance from the centroid of European samples in PC space (x-axis). Vertical black lines on top of the bars in panels (**B,D,F**) show analytical standard errors for the *R*^2^ metric and empirical standard errors for the MSE metric.

On HDL cholesterol, our analysis revealed that one of the main detected drivers of heterogeneity is the participant’s sex (**Figure** 5C; **Figure** S9). After aggregating the mixing proportions by ancestry and sex, we observed that, in addition to the ancestry-matched scores, male and female participants were assigned distinct mixture profiles (**Figure** 5C). This pattern replicated in our analysis of the CARTaGENE biobank, though the differences there are less stark (**Figure** S10; **Figure** S11). This pattern is surprising, however, since the prediction accuracy of the ancestry-stratified scores, quantified in terms of incremental *R*^2^, are nearly identical for males and females (**Figure** 5D). To explain this, we note that `MoEPRS` is not optimizing scale-invariant measures like *R*^2^ and the default version of the model is attuned to heteroscedasticity, or major differences in phenotypic variance across cohort strata. In the UKB dataset, males exhibit ≈ 35% lower variance in HDL cholesterol levels than females, which leads to distinct squared error profiles (**Figure** 5D; **Figure** S11). Thus, partitioning the cohort across sex is meant to reflect this variability, which might be important to highlight for some downstream applications. We also tested a version of `MoEPRS` that partitions the data purely based on the error profiles for each sample, regardless of dispersion patterns (**Methods**). This model no longer identifies distinct profiles for each sex and mostly assigns individuals their ancestry-matched polygenic scores (**Figure** S12; **Figure** S13; see **Discussion**).

Finally, for triglycerides, one of the most visually-discernible trends identified in our analysis is the inverse proportionality between the mixing weight assigned to the European-derived PGS (`EUR`) and distance from the centroid of the European samples in PC space (**Figure** 5C). This effect has been amply documented in systematic analyses of PRS portability [6,7], and our model recapitulated previously observed trends based solely on the error profiles of the ancestry-stratified PRSs (**Figure** 5F). This particular instance shows that when genetic distance to the training cohorts is the main driver of heterogeneity in PRS accuracy, our model is able to pick up on this signal and replicate the behavior of models that are specifically set up to do this task more narrowly [30, 31].

## 4 Discussion

In this work, we presented a flexible modeling framework for ensemble learning of polygenic scores with the aims of improving prediction accuracy on heterogeneous, biobank-scale data while offering researchers practical tools to automatically identify and explore the underlying sources of variability in prediction accuracy. The main benefit of this framework is that, given a set of stratified polygenic scores for a phenotype of interest, it provides personalized, phenotype-specific, and data-driven mixing weights, which should make it easier to deploy these predictors in applied settings. Our experiments on the UK and CARTaGENE biobanks [39, 40] demonstrated that this framework yields modest improvements in prediction accuracy and important insights about how the accuracy of polygenic scores can vary continuously or sharply with age, sex, genetic distance from the training cohort [6, 7, 18].

With these benefits mind, we also highlight a number of limitations to the `MoEPRS` framework as presented here. First, due to the increase in the number of tuning parameters, the model is prone to overfitting, especially on smaller validation cohorts or small sub-cohorts in a larger dataset [38]. Paired with the non-convex objective, this makes the `MoEPRS` model more difficult to train than the `MultiPRS`. Empirically, we observed that while the gating model provided interpretable mixing weights that were relatively stable across biobanks, the mixed scores yielded modest or no gains in prediction accuracy for minority populations. This was disappointing, since improving performance on underrepresented groups was one of the main initial goals for this work. Minority populations in general face two statistical disadvantages: the ancestry-matched polygenic scores are based on smaller training cohorts, and are thus more noisy, and the training sample sizes for the ensemble methods are even more limited. Our interpretation is that the gating model was able to identify the better matching scores, but that higher variance and slight overfitting in these populations might reduced the benefit. In sex-specific analyses, where comparable sample sizes are available, the performance of `MoEPRS` was more consistent and reliable. In short, while our proposed method does reasonably well given the limited sample sizes for minority populations, we anticipate that its performance may be improved further with better representation of minority ancestries as well as more sophisticated training objectives that incorporate regularization techniques [27, 38].

On the conceptual side, there are some caveats to consider when training the `MoEPRS` model on biobank-scale data. One of the main considerations is how to properly account for phenotypic shifts and allele frequency differences across various strata. In our proposed framework, we corrected for these effects by including global covariates into the model [41], which helped steer the gating mechanism towards variation linked to linear genetic effects, rather than spurious environmental influences. As a further consideration, our analyses of the blood biomarker traits revealed that differences in phenotypic variance, or heteroscedasticity [70], can have substantial effects on the behavior of the gating mechanism. Therefore, we caution against misinterpreting the partitions identified by the model as necessarily reflecting differences in the underlying genetic pathways. To ensure that the soft partitions identified in our analyses are robust, we conducted some ablation experiments and quantified the concordance between the mixing patterns produced by applying the model to two independent biobanks [39, 40].

Another consideration for training the `MoEPRS` model is the question of what covariates to include as input to the gating mechanism. In our experiments, we included variables associated with the most explored axes of variation: age, sex, and genomic PCs [5, 11, 17, 18]. For more comprehensive analyses of specific phenotypes, researchers might consider including other variables of interest based on epidemiological evidence. In general, if the true drivers of heterogeneity are not represented in the gating model input features, then proxy features may be highlighted, as in the case of using age and sex to triangulate samples that are likely to take cholesterol-lowering medication.

Despite these current limitations, we believe that the `MoEPRS` framework is flexible and foresee that it might be applied and extended along a number of promising directions. One major direction that may yield tangible benefits is to update the gating mechanism to use non-linear functions of the covariates, such as deep neural networks [71]. These networks can then be trained in a multi-task manner [72], across related phenotypes and diverse biobanks, to optimize prediction accuracy and reduce overfitting. Furthermore, this setup can be naturally paired with a pre-training/fine-tuning paradigm [73] to transfer learned mappings from large-scale biobanks to smaller test cohorts, e.g. samples recruited for a specific clinical trial. The second major extension that we anticipate is reconfiguring the model to learn different mixing proportions for each genomic segment separately, which might greatly improve its performance on samples with recent genetic admixture [74, 75]. Finally, recognizing that the model provides valuable tools for uncovering heterogeneity, one potential avenue of future research is to use the model for predicting disease phenotypes using polygenic scores for related [26, 27] or endophenotypes [76], which can help uncover disease subtypes in a data-driven manner [77].

## Code Availability

- Scripts to replicate all the analyses and generate the figures for this manuscript are available via GitHub: https://github.com/shz9/moe-prs-paper
- `viprs` software used to infer polygenic scores for sex-differentiated phenotypes: https://github.com/shz9/viprs
- `pgsc_calc` software utility for computing polygenic scores and continental ancestry prediction: https://github.com/PGScatalog/pgsc_calc

## Data Availability

- GWAS summary statistics for sex-differentiated phenotypes: https://zenodo.org/records/7222725

## Acknowledgements

We thank members of the Gravel and Li labs, Marc-André Legault, and Celia Greenwood for useful feedback and discussions on earlier drafts of this manuscript. We are grateful to Alex Diaz-Papkovich for providing the UMAP+HDBSCAN clusters used in some of the analyses and to Zhe Xie and Tim D’Aoust for testing and experimenting with earlier versions of the `MoEPRS` software. This research used the NeuroHub infrastructure and was undertaken thanks in part to funding from the Canada First Research Excellence Fund, awarded through the Healthy Brains, Healthy Lives initiative at McGill University. This research was enabled in part by support provided by Calcul Québec and the Digital Research Alliance of Canada. S.G. was supported by the Canadian Institute for Health Research (CIHR) project grant 437576, NSERC grant (RGPIN-2017-04816), the Canada Research Chair program, and the Canada Foundation for Innovation. Y.L. was supported by the following funding programs: the Canada Research Chair (Tier 2) in Machine Learning for Genomics and Healthcare (CRC-2021-00547), Natural Sciences and Engineering Research Council (NSERC) Discovery Grant (RGPIN-2016-05174).

## Author contributions

S.Z., S.G. and Y.L. conceived the study. S.Z. derived the initial formulation of the `MoEPRS` model. S.G. and Y.L. provided input and feedback to arrive at the final version of the model presented here. S.Z. conducted UKB and CaG experiments and generated the main figures and results. S.Z. wrote the initial manuscript. All authors edited and reviewed the final version. S.G. and Y.L. supervised the work.

