## Supplementary for "Personalized polygenic risk prediction and assessment with a Mixture-of-Experts framework"

### S1 Supplementary Figures

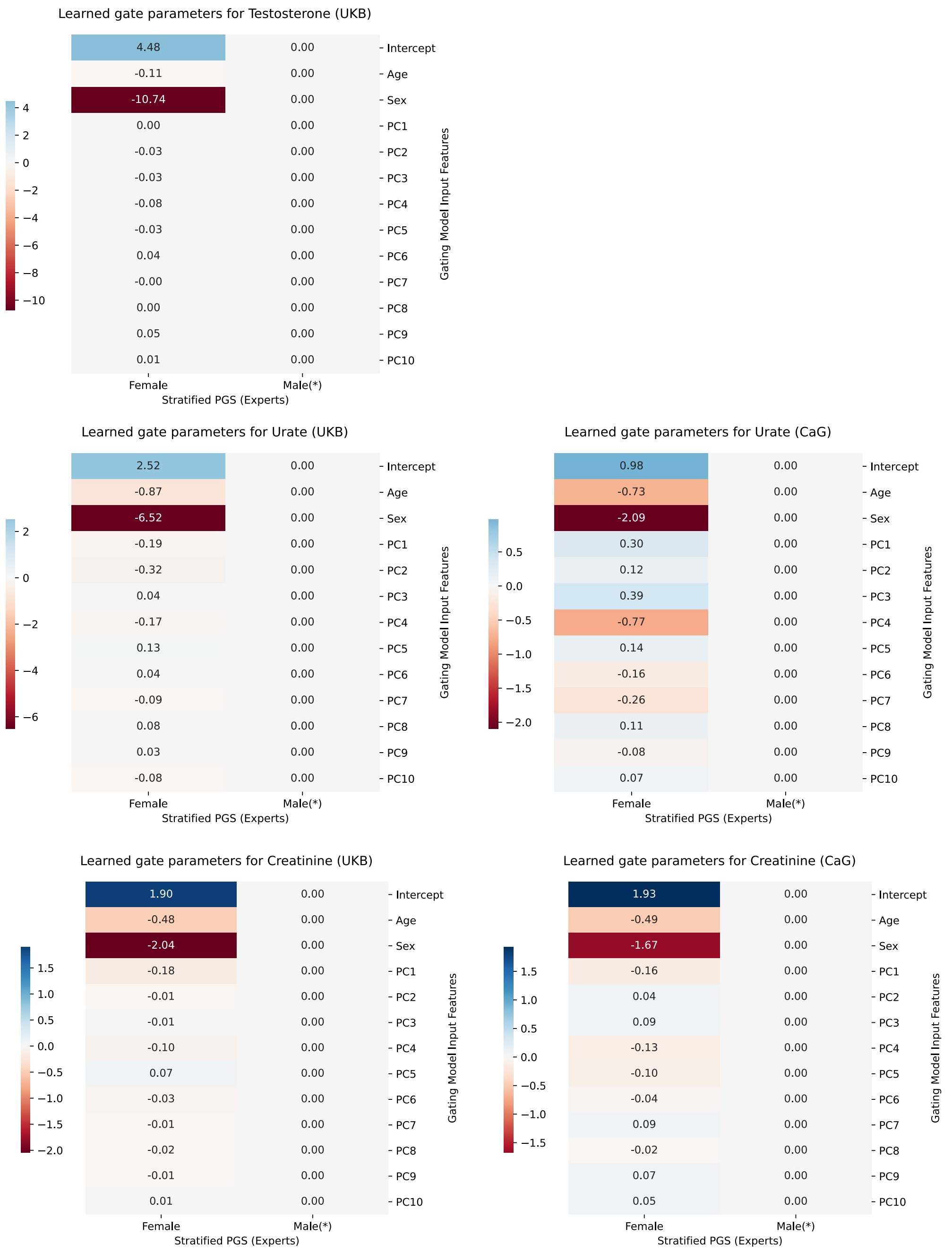


Figure S1: Heatmap showing the gating model parameters inferred from fitting `MoEPRS` on sex-differentiated blood biomarker traits in the UK (left) and CARTaGENE (right) biobanks. The first row shows the inferred gating model parameters for testosterone, the second row shows model parameters for urate, and the third row shows parameters for creatinine. The y-axis shows the covariates used as input to the gating model and x-axis shows the single source PGSs that the gating model learned to mix on each cohort (**Supplementary Table** S1). The PGS denoted with (*) is used as a reference class and its parameters are zeroed out to ensure identifiability. Each cell shows the parameter value and the background color indicates its magnitude and direction.

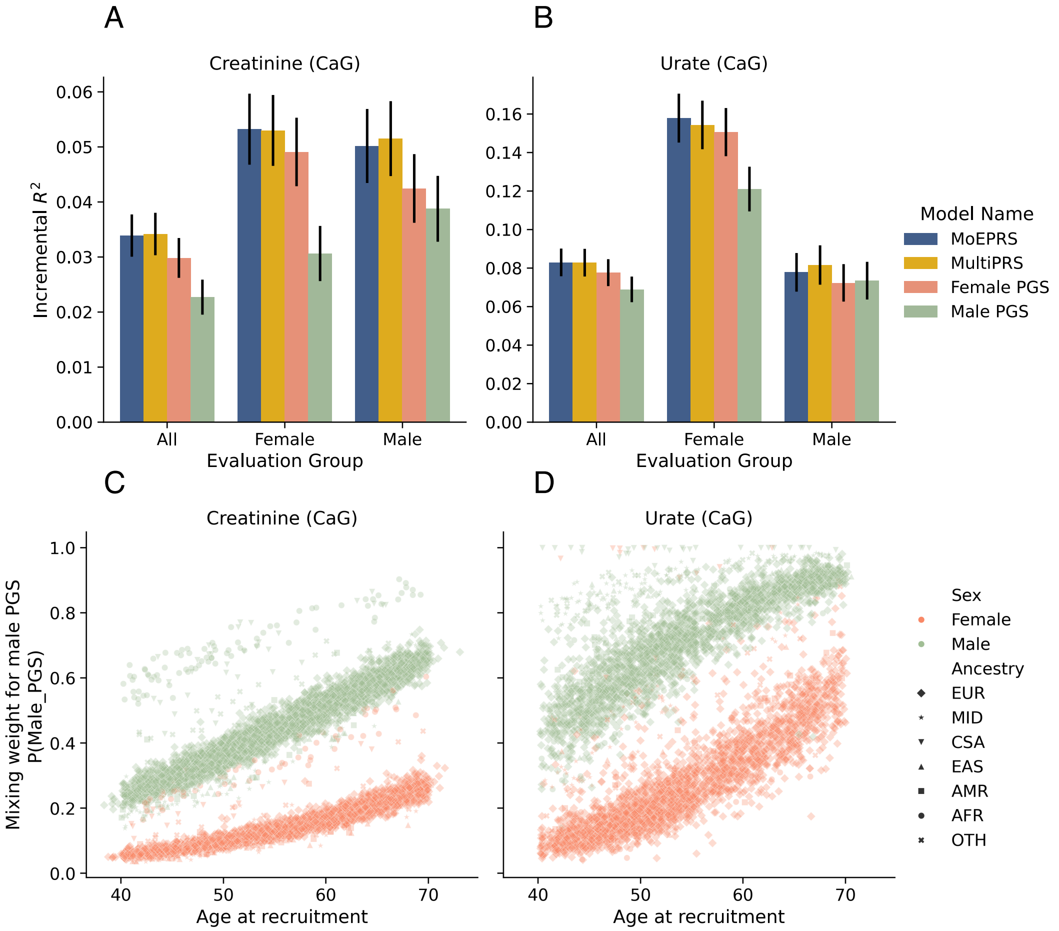


Figure S2: Prediction accuracy and specialization behavior of the `MoEPRS` model on sex-differentiated complex traits in the CARTaGENE (CaG) biobank. Sex-stratified polygenic scores ("Male PGS" and "Female PGS") derived from European samples in the UK Biobank were used as input to predict phenotypes for the entire CaG cohort. Ensemble PRS models that combine the stratified scores include our proposed `MoEPRS` as well as the popular baseline `MultiPRS`, which learns a linear combination of individual scores. The complex traits analyzed are Creatinine and Urate levels measured in blood biochemistry assays. Panels (**A-B**) show prediction accuracy in terms of incremental $R^2$ on the held-out test set in CaG. The accuracy measures are stratified by sex (x-axis), and vertical black lines on top of the bars show analytical standard errors for the $R^2$ metric. Panels (**C-D**) show the weight assigned to the "Male PGS" by the gating model for each individual in the test set, as a function of the sample's age at recruitment (x-axis). The scatter points are differentiated by colors (green for males and orange for females) and markers corresponding to different continental ancestry groups. The ancestry groups include: `EUR` (European), `MID` (Middle Eastern), `CSA` (Central and South Asian), `EAS` (East Asian), `AMR` (Admixed American), `AFR` (African), and `OTH` (Other).

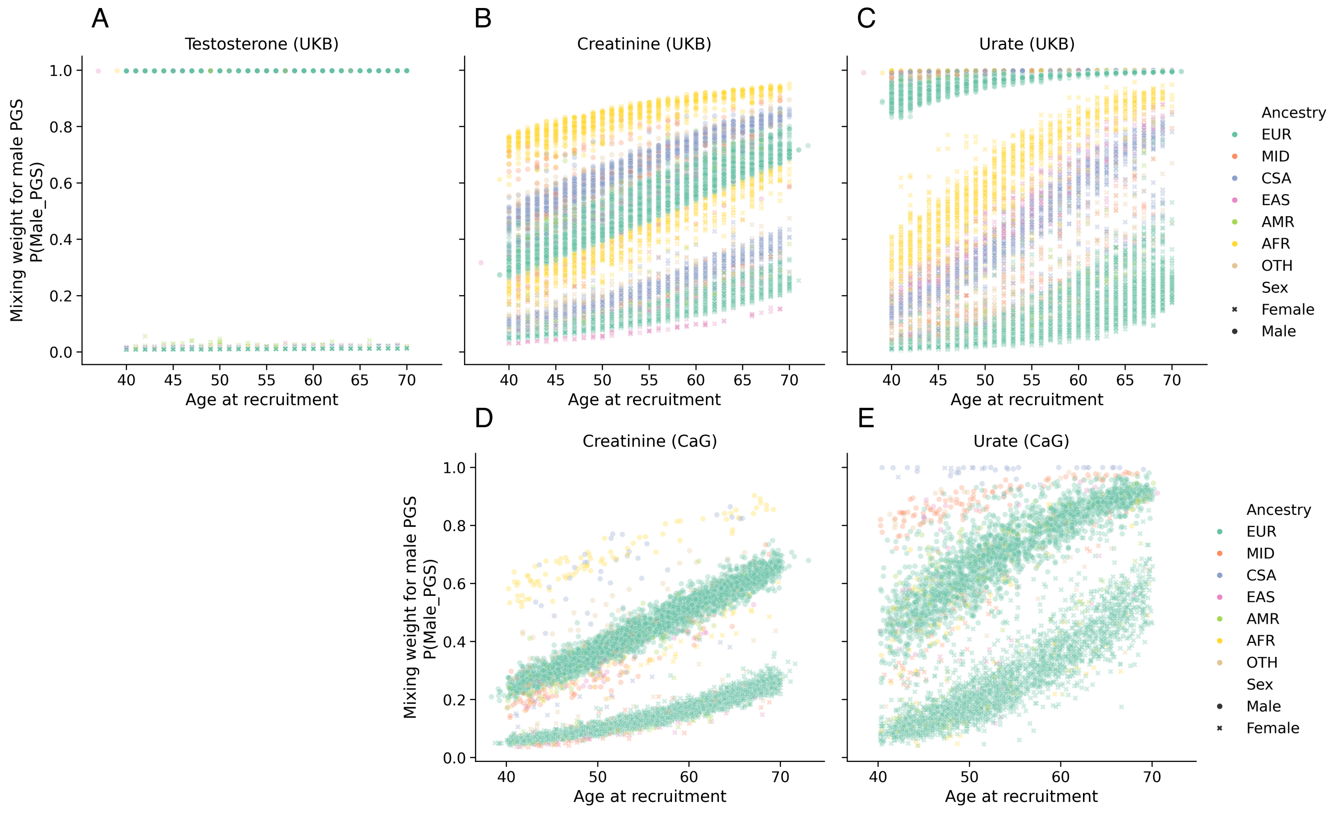


Figure S3: Specialization behavior of the `MoEPRS` model on sex-differentiated complex traits in the CARTaGENE (CaG) and UK (UKB) biobanks. Each panel shows the weight assigned to the "Male PGS" by the gating model for each individual in the respective test set, as a function of the sample's age at recruitment (x-axis). The gating models were fitted to the training data from each biobank separately and independently. The scatter points are differentiated by markers (• for males and × for females) and colors corresponding to different continental ancestry groups. The ancestry groups include: `EUR` (European), `MID` (Middle Eastern), `CSA` (Central and South Asian), `EAS` (East Asian), `AMR` (Admixed American), `AFR` (African), and `OTH` (Other).

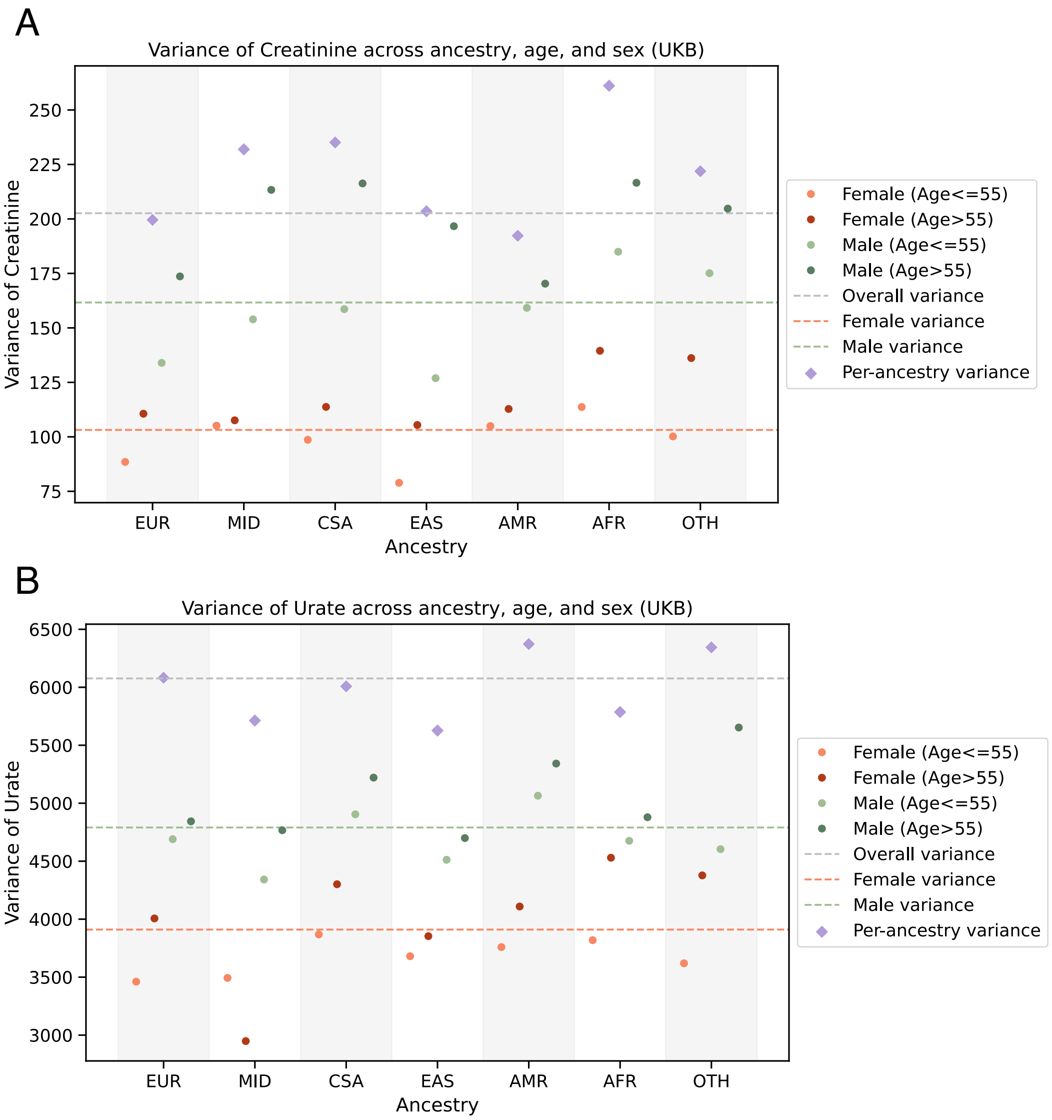


Figure S4: Phenotypic variance of sex-differentiated phenotypes in the UK Biobank, stratified by ancestry, age and sex. The phenotypes included are creatinine (**A**) and urate (**B**) levels measured in blood biochemistry assays. The x-axis shows predicted ancestry group for the sample of individuals and the y-axis records the variance of the phenotype in each stratum. The samples are further differentiated by sex and age groups. Dashed lines show the overall phenotypic variance (silver), female variance (orange), and male variance (green).

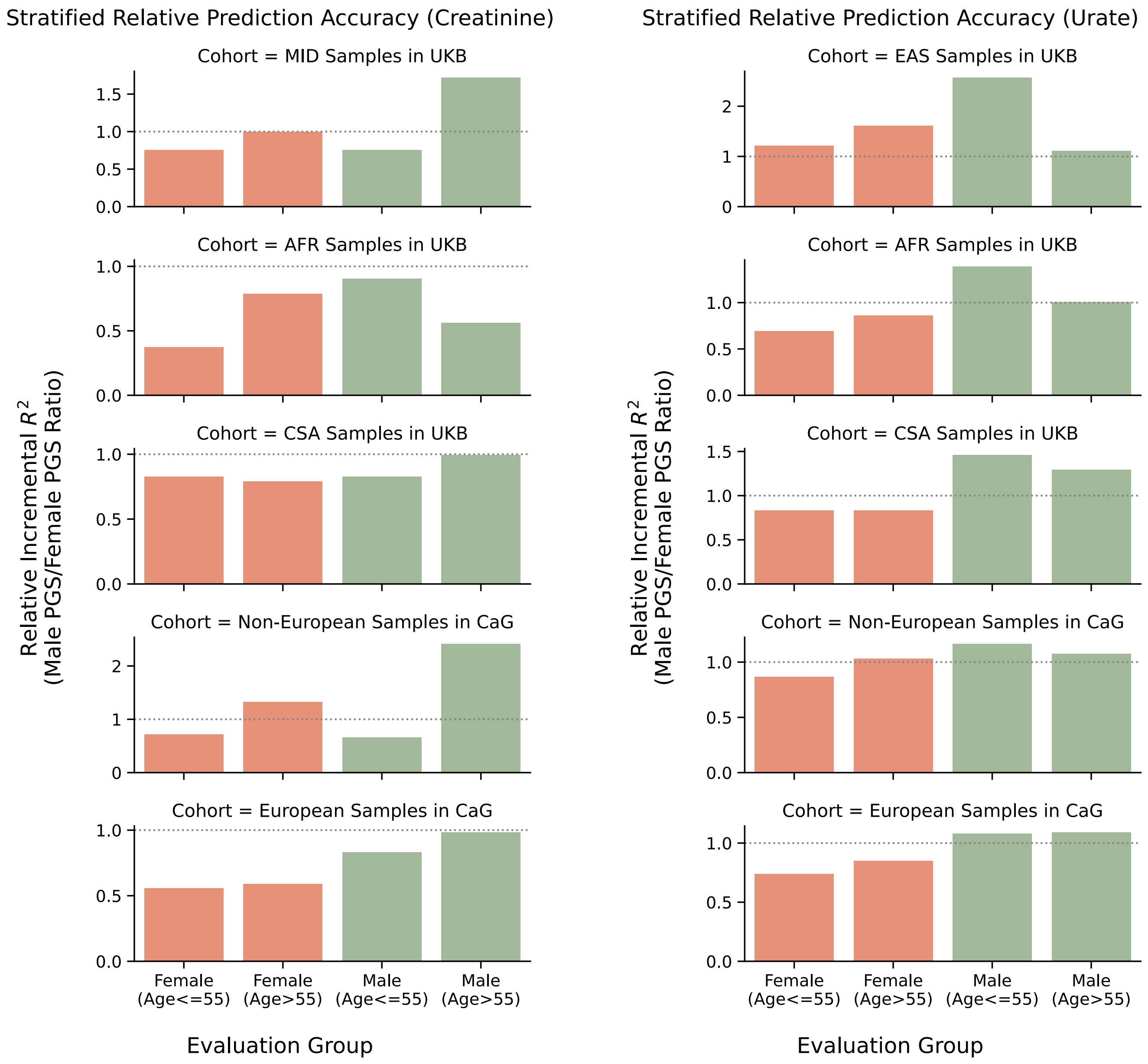


Figure S5: Relative prediction accuracy of the sex-stratified polygenic scores across different sub-cohorts in the CARTaGENE (CaG) and UK (UKB) biobanks. Each subpanel compares the performance of male-derived ("Male PGS") to the female-derived ("Female PGS") on a particular sub-cohort by examining the ratio of their incremental $R^2$ metric. Left/right panels show relative performance in predicting Creatinine/Urate levels in the blood. The sub-cohorts include different ancestry groups (as specified in the title of each sub-panel) as well as sex and age groups (x-axis). The specific ancestry groups examined include `AFR` (Africans), `MID` (Middle Eastern), `EAS` (East Asian), and `CSA` (Central and South Asian). Male sub-cohorts are colored in green while female sub-cohort are colored in orange. This figure shows that the relative performance of the sex-stratified PGSs is not uniform and varies by sex, age, and ancestry.

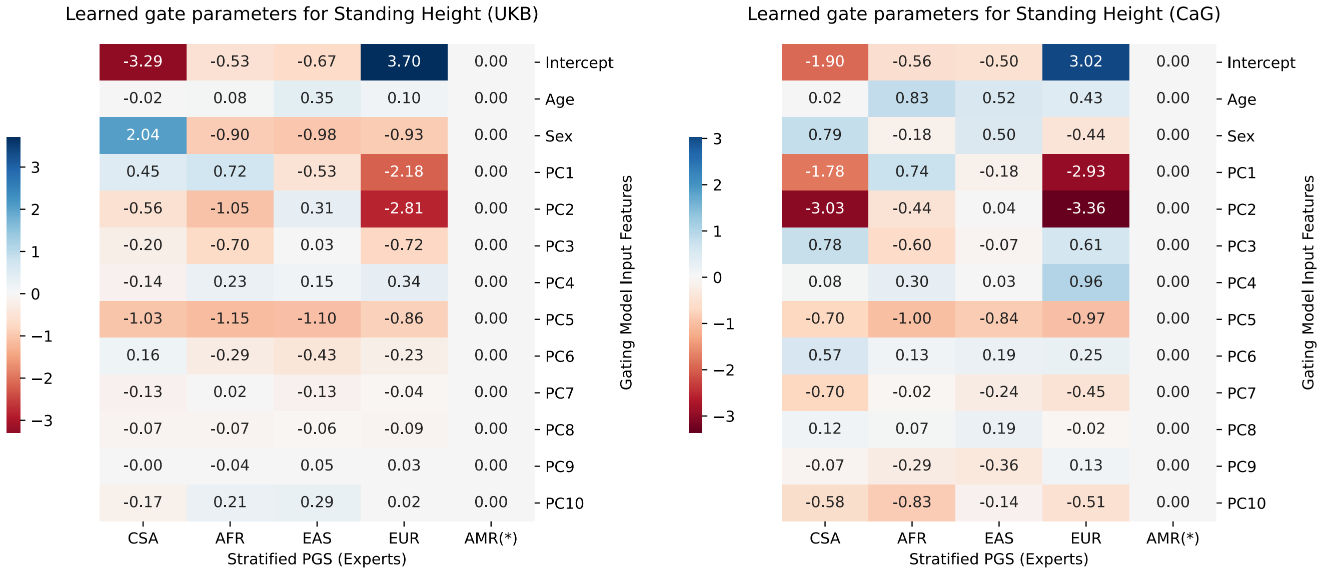


| UKB | CSA | AFR | EAS | EUR | AMR(*) |
|---|---|---|---|---|---|
| Intercept | -3.29 | -0.53 | -0.67 | 3.70 | 0.00 |
| Age | -0.02 | 0.08 | 0.35 | 0.10 | 0.00 |
| Sex | 2.04 | -0.90 | -0.98 | -0.93 | 0.00 |
| PC1 | 0.45 | 0.72 | -0.53 | -2.18 | 0.00 |
| PC2 | -0.56 | -1.05 | 0.31 | -2.81 | 0.00 |
| PC3 | -0.20 | -0.70 | 0.03 | -0.72 | 0.00 |
| PC4 | -0.14 | 0.23 | 0.15 | 0.34 | 0.00 |
| PC5 | -1.03 | -1.15 | -1.10 | -0.86 | 0.00 |
| PC6 | 0.16 | -0.29 | -0.43 | -0.23 | 0.00 |
| PC7 | -0.13 | 0.02 | -0.13 | -0.04 | 0.00 |
| PC8 | -0.07 | -0.07 | -0.06 | -0.09 | 0.00 |
| PC9 | -0.00 | -0.04 | 0.05 | 0.03 | 0.00 |
| PC10 | -0.17 | 0.21 | 0.29 | 0.02 | 0.00 |

| CaG | CSA | AFR | EAS | EUR | AMR(*) |
|---|---|---|---|---|---|
| Intercept | -1.90 | -0.56 | -0.50 | 3.02 | 0.00 |
| Age | 0.02 | 0.83 | 0.52 | 0.43 | 0.00 |
| Sex | 0.79 | -0.18 | 0.50 | -0.44 | 0.00 |
| PC1 | -1.78 | 0.74 | -0.18 | -2.93 | 0.00 |
| PC2 | -3.03 | -0.44 | 0.04 | -3.36 | 0.00 |
| PC3 | 0.78 | -0.60 | -0.07 | 0.61 | 0.00 |
| PC4 | 0.08 | 0.30 | 0.03 | 0.96 | 0.00 |
| PC5 | -0.70 | -1.00 | -0.84 | -0.97 | 0.00 |
| PC6 | 0.57 | 0.13 | 0.19 | 0.25 | 0.00 |
| PC7 | -0.70 | -0.02 | -0.24 | -0.45 | 0.00 |
| PC8 | 0.12 | 0.07 | 0.19 | -0.02 | 0.00 |
| PC9 | -0.07 | -0.29 | -0.36 | 0.13 | 0.00 |
| PC10 | -0.58 | -0.83 | -0.14 | -0.51 | 0.00 |

Figure S6: Heatmap showing the gating model parameters inferred from fitting `MoEPRS` on Standing Height data in the UK (left) and CARTaGENE (right) biobanks. The y-axis shows the covariates used as input to the gating model and x-axis shows the single source height PGSs that the gating model learned to mix on each cohort (**Supplementary Table** S1). The PGS denoted with (*) is used as a reference class and its parameters are zeroed out to ensure identifiability. Each cell shows the parameter value and the background color indicates its magnitude and direction.

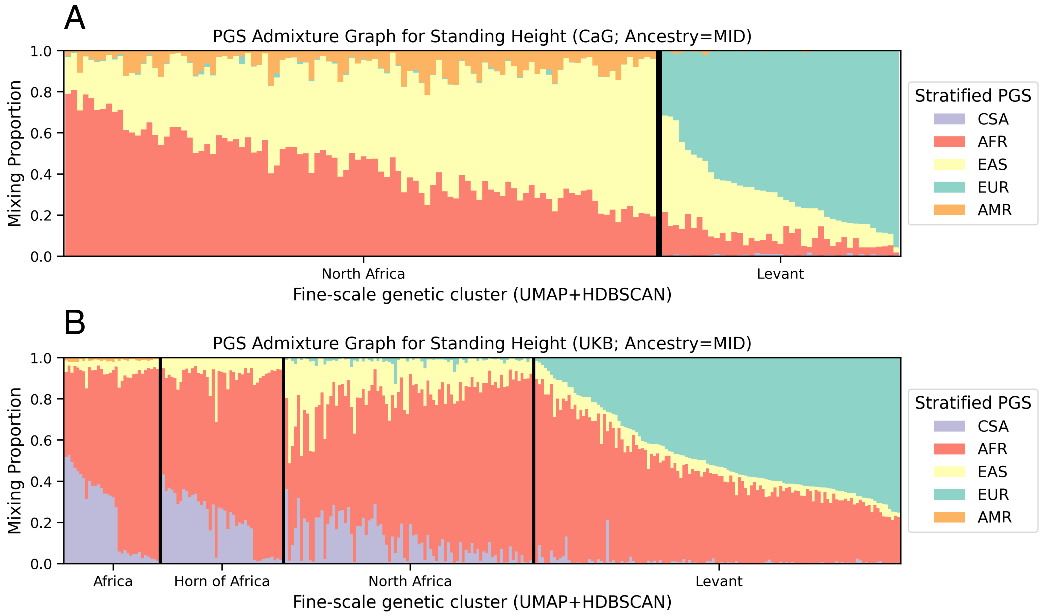


Figure S7: Specialization behavior of the `MoEPRS` model in predicting Standing Height in samples of "Middle Eastern" (MID) ancestry in the UK (UKB) and CARTaGENE (CaG) biobanks. Both panels show fine-grained PGS admixture graphs, which represent the mixing weights assigned by the gating model for individuals in the test set, stratified by their assigned UMAP cluster. Colors correspond to one of the ancestry-stratified PGS models for height (PGS Catalog Publication ID: `PGP000382`): Central and South Asian PGS (CSA; purple), African PGS (AFR; red), East Asian PGS (EAS; yellow), European PGS (EUR; turquoise), and Admixed American PGS (AMR; orange). The UMAP+HDBSCAN clusters in CARTaGENE are `4-NAF` (North African), `5-MIE` (Middle Eastern), the latter is mostly individuals of Lebanese origin. The UMAP clusters in the UK Biobank are `18 AFR` (Sub-Saharan African), `23 HAFR` (Horn of Africa), `5 LEV` (Levantine ancestry), and `6 NAF` (North African). This figure shows that when trained on data for the height phenotype on each biobank separately, the model does not treat "continental ancestry" as a monolith and finds differential and replicated patterns for the sub-populations present.

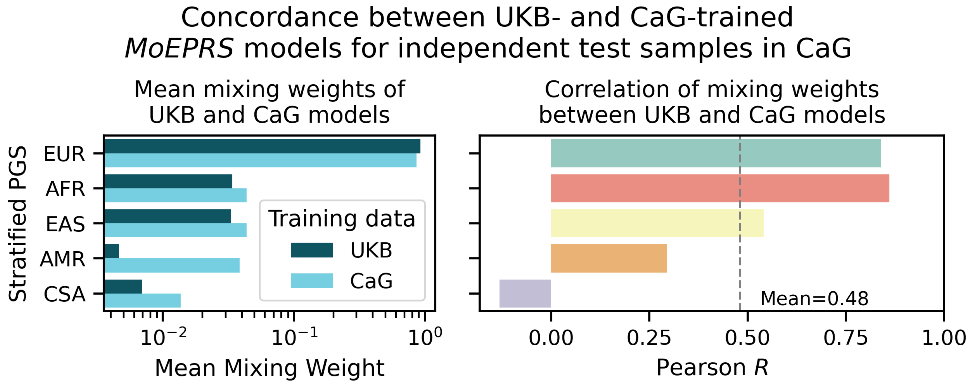


Figure S8: Concordance between mixing weights assigned by `MoEPRS` models trained on Standing Height data in two different biobanks. After the gating models were trained on the training subset in the UK (UKB) and CARTaGENE (CaG) biobanks, they were applied to independent test samples in CARTaGENE. The figure shows quantitatively the level of agreement between the mixing weights assigned by the two models, both in terms of overall proportions (left) and how well they correlate for each PGS individually (right). Dashed vertical line in the right sub-plot and accompanying text annotation show the mean correlation of the mixing weights across all the five models.

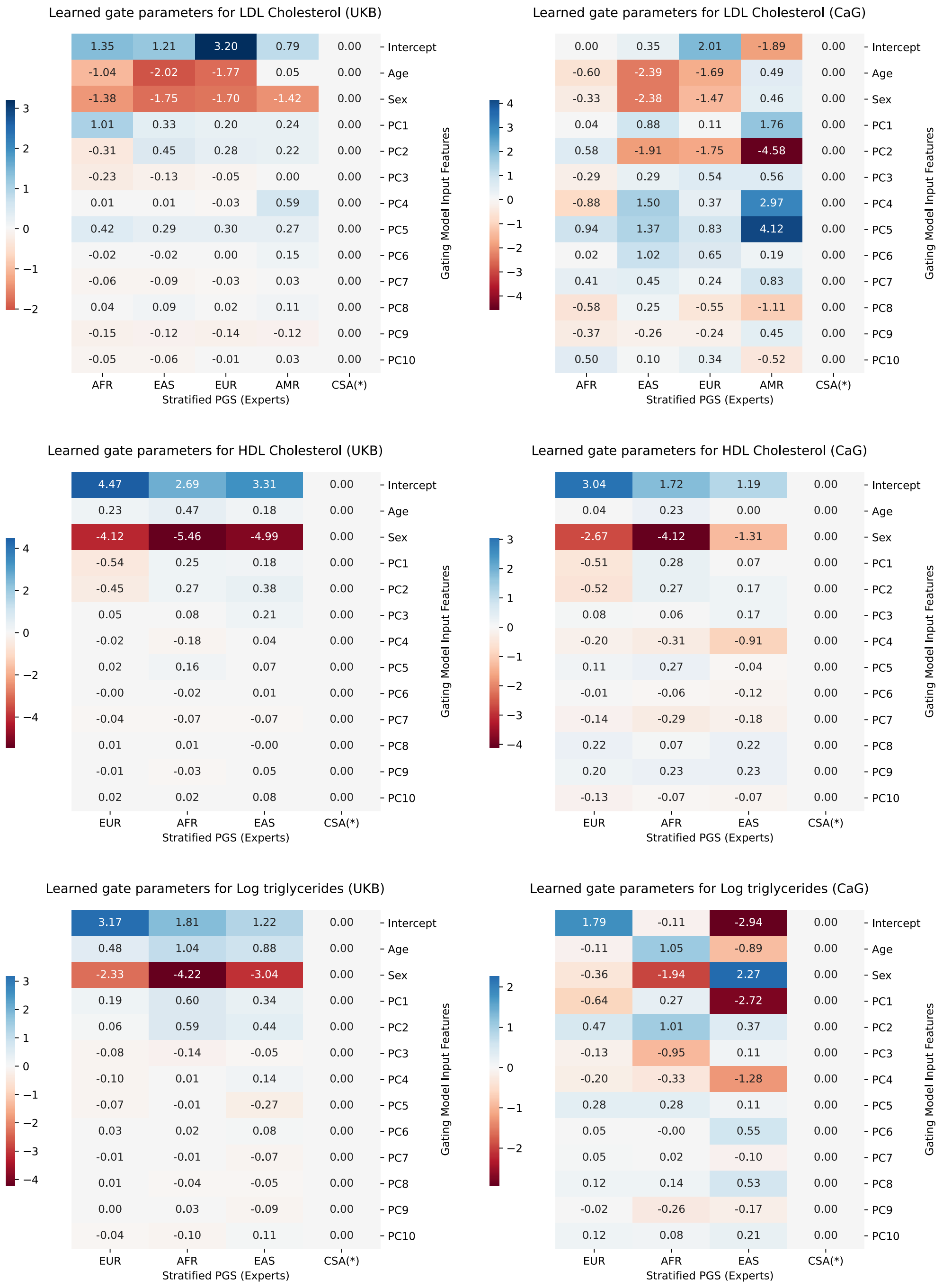


Figure S9: Heatmap showing the gating model parameters inferred from fitting `MoEPRS` on blood lipid traits in the UK (left) and CARTaGENE (right) biobanks. The first row shows the inferred gating model parameters for LDL cholesterol, the second row shows model parameters for HDL cholesterol, and the third row shows parameters for log(triglycerides). The y-axis shows the covariates used as input to the gating model and x-axis shows the single source PGSs that the gating model learned to mix on each cohort (**Supplementary Table** S1). The PGS denoted with `(*)` is used as a reference class and its parameters are zeroed out to ensure identifiability. Each cell shows the parameter value and the background color indicates its magnitude and direction.

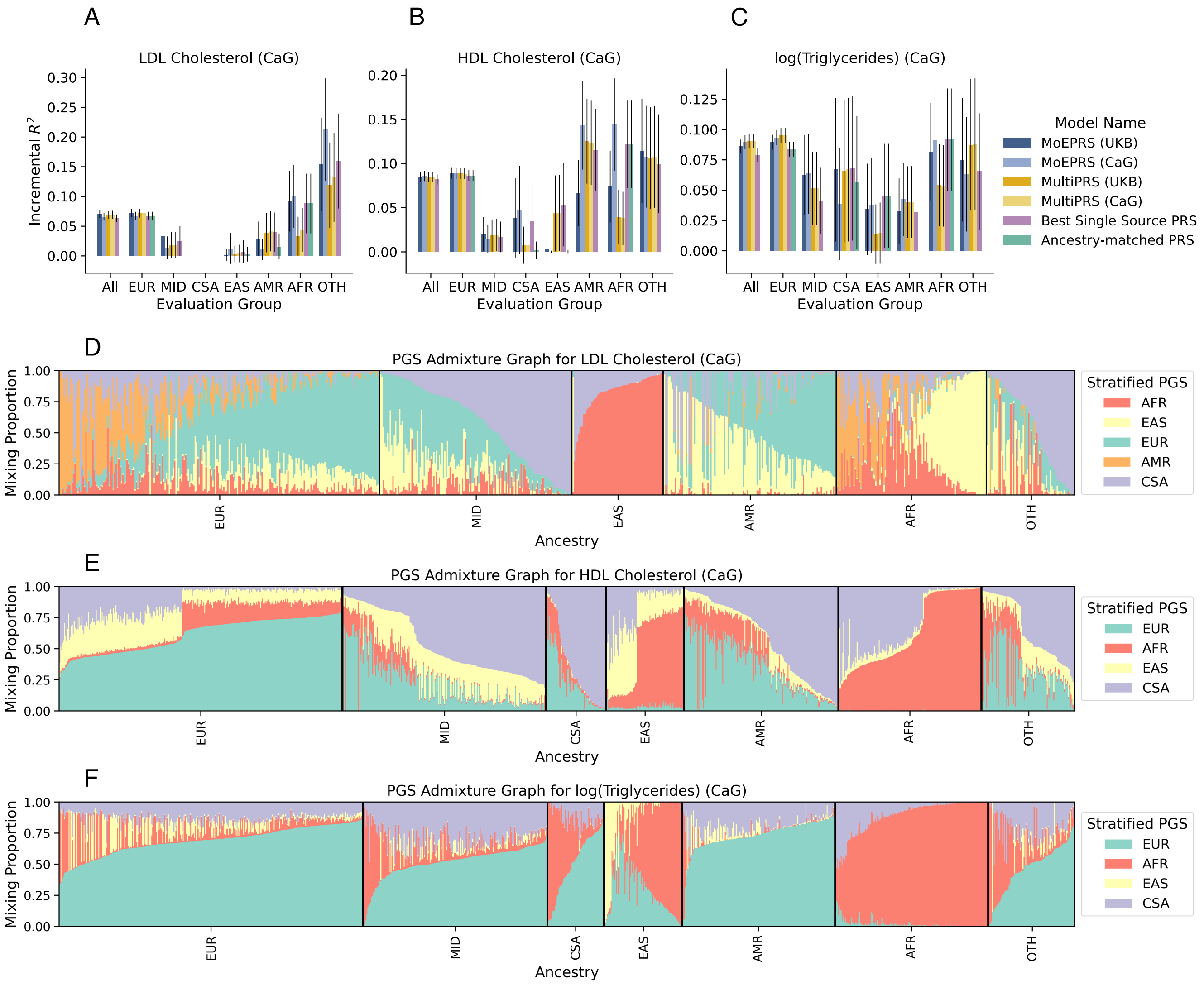


Figure S10: Prediction accuracy and specialization behavior of the `MoEPRS` model on blood lipid traits in the CARTaGENE biobank (CaG). Trait-specific and ancestry-stratified polygenic scores derived from up to five continental ancestry groups were used as input to predict each molecular phenotype. The traits analyzed are low-density lipoprotein (LDL) cholesterol, high-density lipoprotein (HDL) cholesterol, and log-transformed triglyceride levels in the blood. Ensemble PRS models that combine the stratified scores include our proposed `MoEPRS` as well as the popular baseline `MultiPRS`, which learns a linear combination of individual scores. Panels (**A-C**) show prediction accuracy in terms of incremental $R^2$ on the held-out test set in CaG. The accuracy measures are stratified by ancestry (x-axis), and vertical black lines on top of the bars show analytical standard errors for the $R^2$ metric. Panels (**D-F**) show PGS admixture graphs for each phenotype, which represent the mixing weights assigned by the gating model for a random subsample of individuals, stratified by continental ancestry.

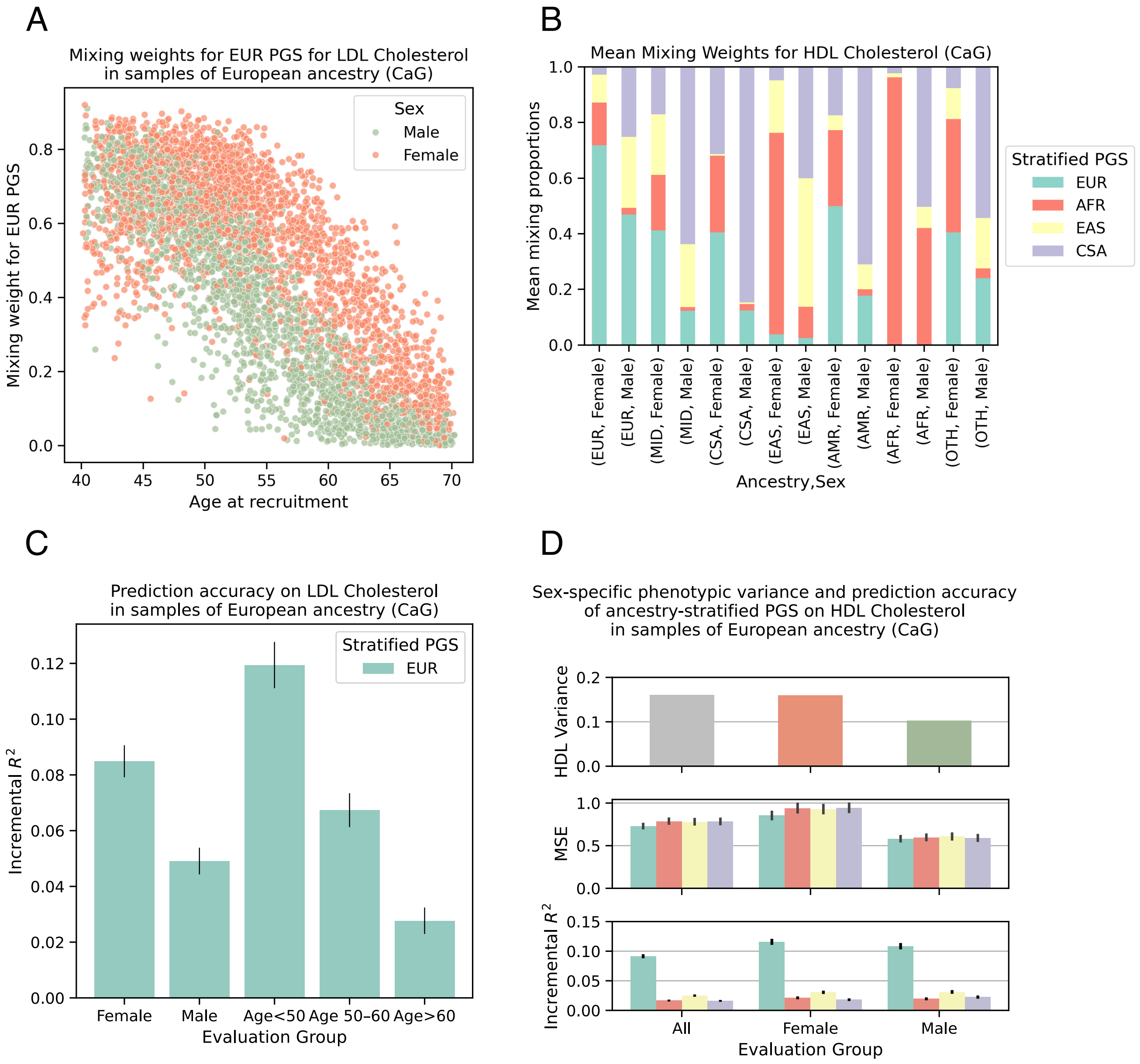


Figure S11: Investigation of some of the soft cohort partitions identified by `MoEPRS` in the analysis of blood lipid phenotypes in the CARTaGENE biobank (CaG). Panel (**A**) shows the mixing weight assigned to the European-derived PGS (`EUR`) model for LDL cholesterol as a function of the sample's age at recruitment (x-axis) and sex (color). Panel (**B**) shows the average mixing proportions assigned by `MoEPRS` to each group of individuals when predicting HDL cholesterol, categorized by the sample's predicted continental ancestry and sex (x-axis). Panel (**C**) shows the prediction accuracy, in terms of incremental $R^2$, of the `EUR` PGS in European samples in the CaG, stratified by sex and age groups (x-axis). Panel (**D**) shows, for each sex, the phenotypic variance of HDL cholesterol as well as the predictive performance of the ancestry-stratified PGSs on European samples in the UKB. The predictive performance metrics include Mean Squared Error (MSE) of the covariates-augmented models as well as Incremental $R^2$. Vertical black lines on top of the bars in panels (**C-D**) show analytical standard errors for the $R^2$ metric.

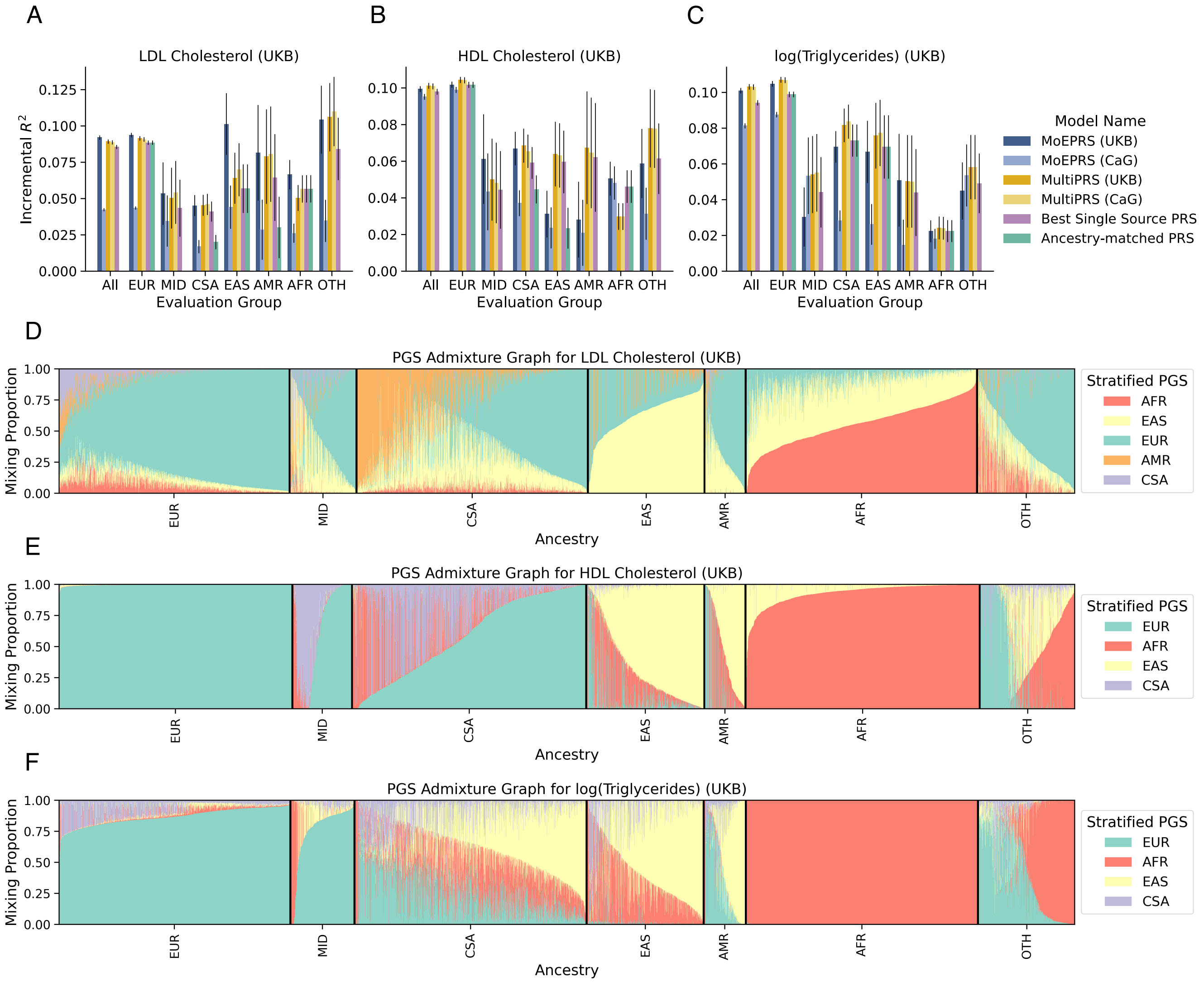


Figure S12: Prediction accuracy and specialization behavior of the MoEPRS model on blood lipid traits in the UK Biobank (UKB). The ensemble model was fit with fixed variance for all the PGS experts ($\sigma_k = 1$), thus only considering squared error patterns as the sole criterion for the gating mechanism. Trait-specific and ancestry-stratified polygenic scores derived from up to five continental ancestry groups were used as input to predict each molecular phenotype. The traits analyzed are low-density lipoprotein (LDL) cholesterol, high-density lipoprotein (HDL) cholesterol, and log-transformed triglyceride levels in the blood. Ensemble PRS models that combine the stratified scores include our proposed MoEPRS as well as the popular baseline MultiPRS, which learns a linear combination of individual scores. Panels (**A-C**) show prediction accuracy in terms of incremental $R^2$ on the held-out test set in the UKB. The accuracy measures are stratified by ancestry (x-axis), and vertical black lines on top of the bars show analytical standard errors for the $R^2$ metric. Panels (**D-F**) show PGS admixture graphs for each phenotype, which represent the mixing weights assigned by the gating model for a random subsample of individuals, stratified by continental ancestry.

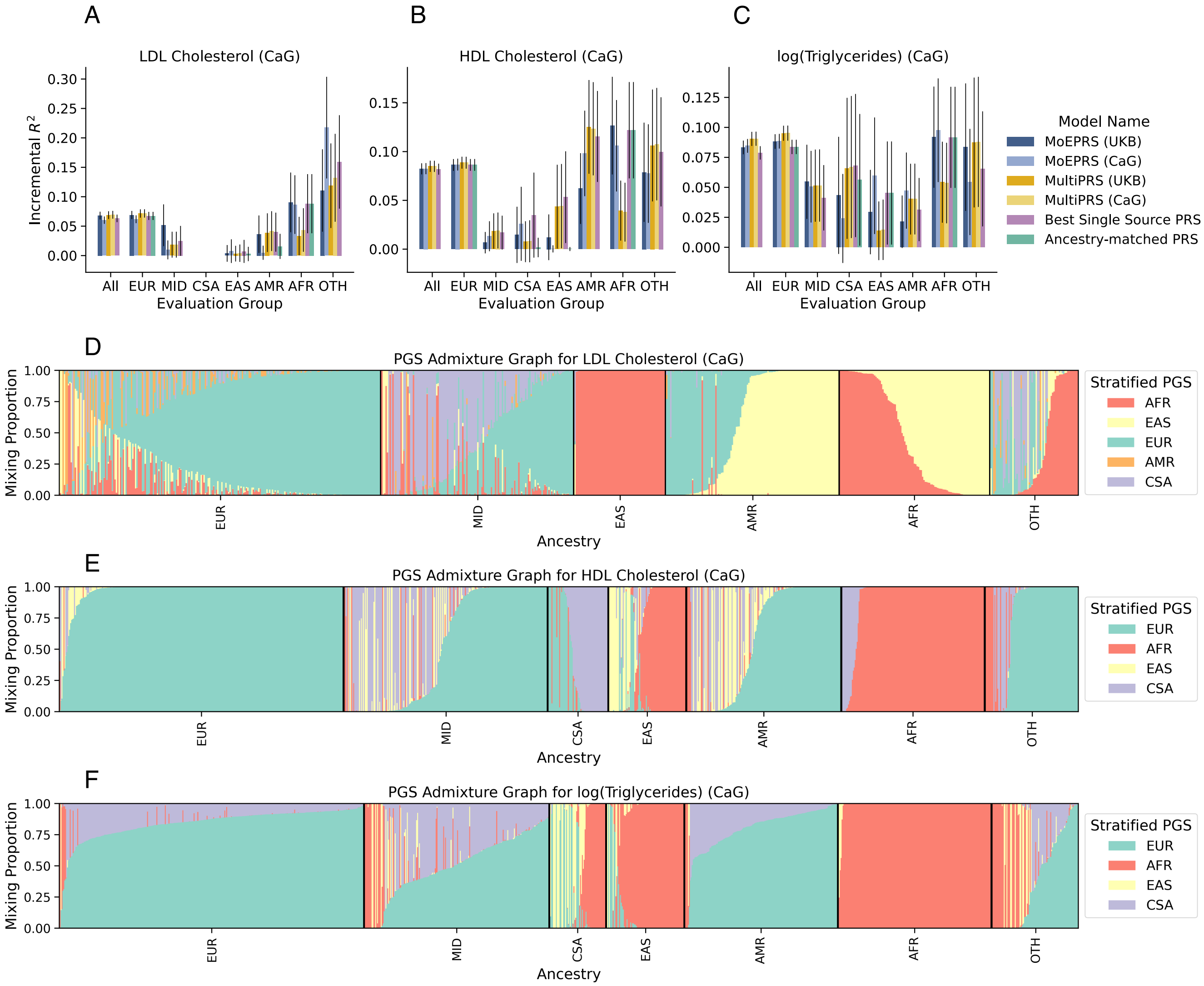


Figure S13: Prediction accuracy and specialization behavior of the MoEPRS model on blood lipid traits in the CARTaGENE biobank (CaG). The ensemble model was fit with fixed variance for all the PGS experts ($\sigma_k = 1$), thus only considering squared error patterns as the sole criterion for the gating mechanism. Trait-specific and ancestry-stratified polygenic scores derived from up to five continental ancestry groups were used as input to predict each molecular phenotype. The traits analyzed are low-density lipoprotein (LDL) cholesterol, high-density lipoprotein (HDL) cholesterol, and log-transformed triglyceride levels in the blood. Ensemble PRS models that combine the stratified scores include our proposed MoEPRS as well as the popular baseline MultiPRS, which learns a linear combination of individual scores. Panels (**A-C**) show prediction accuracy in terms of incremental $R^2$ on the held-out test set in CaG. The accuracy measures are stratified by ancestry (x-axis), and vertical black lines on top of the bars show analytical standard errors for the $R^2$ metric. Panels (**D-F**) show PGS admixture graphs for each phenotype, which represent the mixing weights assigned by the gating model for a random subsample of individuals, stratified by continental ancestry.

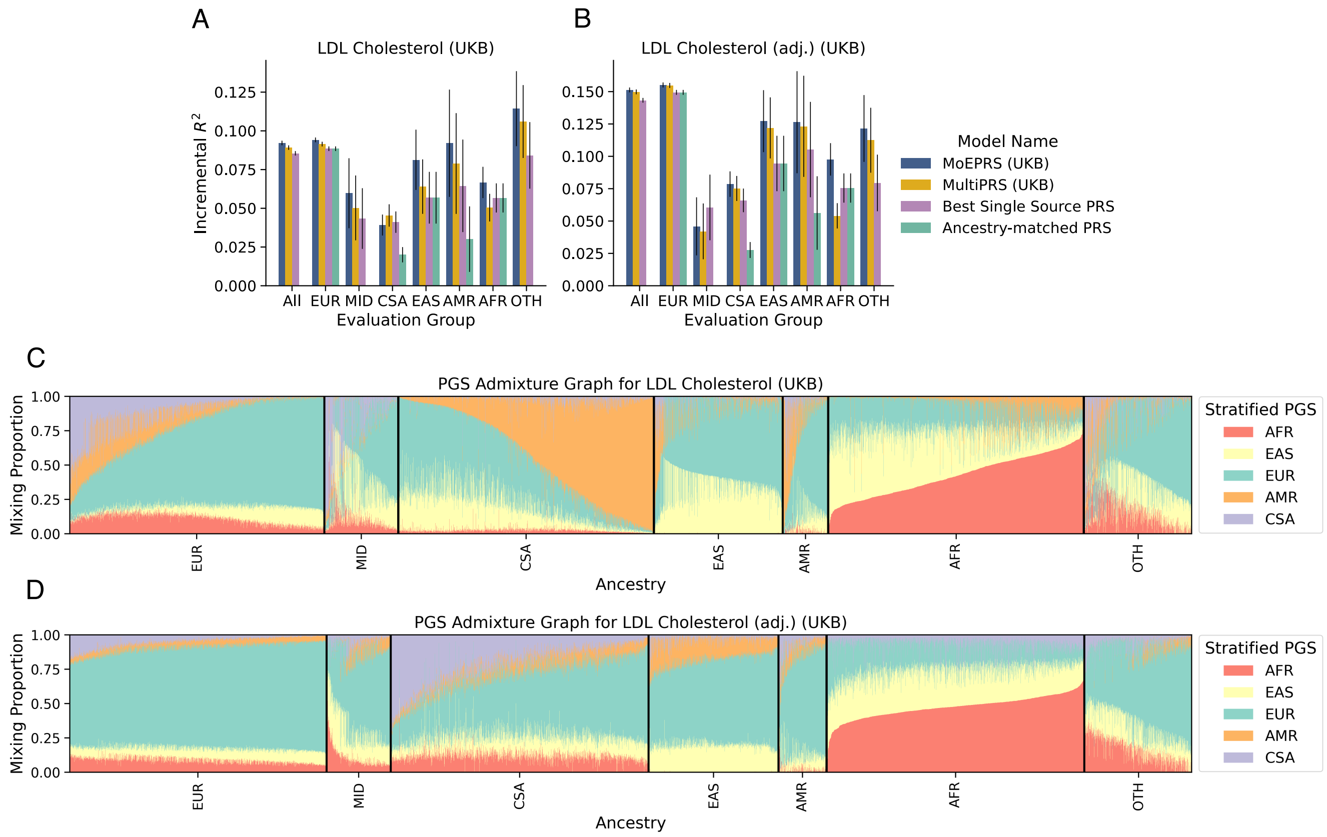


Figure S14: Prediction accuracy and specialization behavior of the `MoEPRS` model on LDL Cholesterol in the UK Biobank (UKB). Trait-specific and ancestry-stratified polygenic scores derived from up to five continental ancestry groups were used as input to predict cholesterol levels. The traits analyzed are direct LDL cholesterol measurements as well as an adjusted LDL cholesterol, where samples who reported using cholesterol-lowering medication had their measured values divided by $0.7$. Ensemble PRS models that combine the stratified scores include our proposed `MoEPRS` as well as the popular baseline `MultiPRS`, which learns a linear combination of individual scores. Panels (**A-B**) show prediction accuracy in terms of incremental $R^2$ on the held-out test set in the UKB. The accuracy measures are stratified by ancestry (x-axis), and vertical black lines on top of the bars show analytical standard errors for the $R^2$ metric. Panels (**C-D**) show PGS admixture graphs for each phenotype, which represent the mixing weights assigned by the gating model for a random subsample of individuals, stratified by continental ancestry.

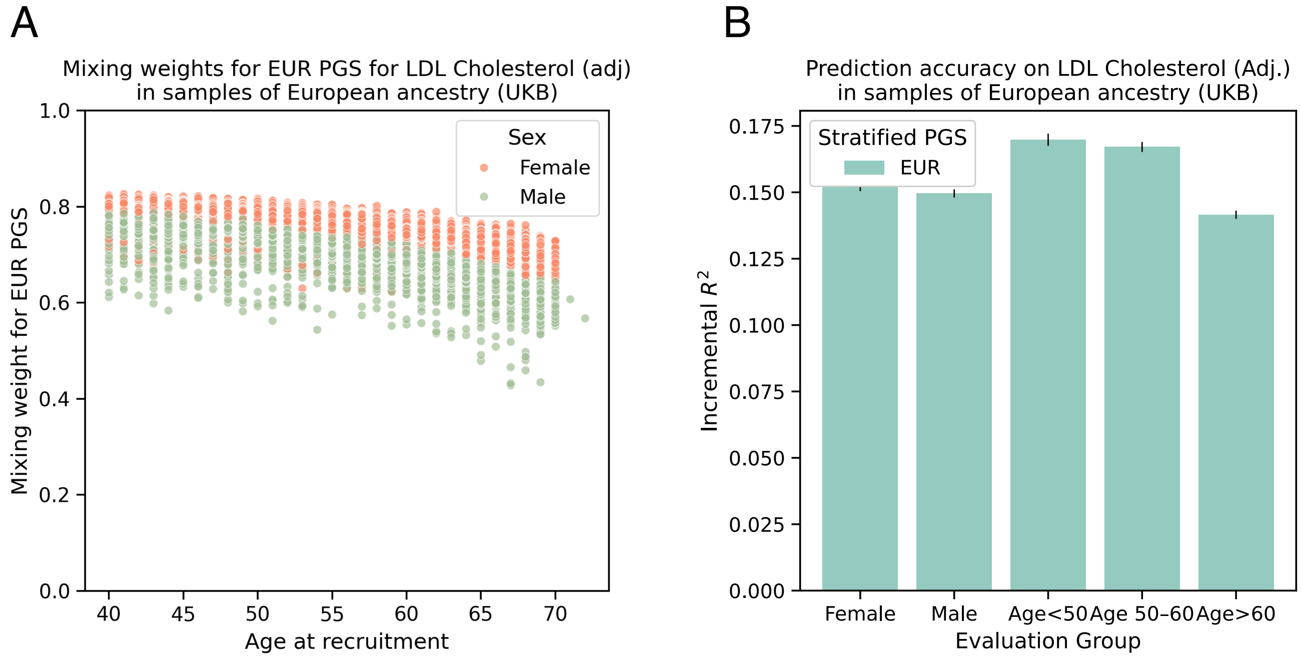


Figure S15: Investigation of the mixing weights identified by `MoEPRS` in the analysis of adjusted LDL Cholesterol phenotype in the UK Biobank. The adjustment for LDL cholesterol consisted of dividing the direct measurement by $0.7$ for where samples who reported using cholesterol-lowering medication. Panel (**A**) shows the mixing weight assigned to the European-derived PGS (`EUR`) model for LDL cholesterol (adj.) as a function of the sample's age at recruitment (x-axis) and sex (color). Panel (**B**) shows the prediction accuracy of the `EUR` PGS for LDL Cholesterol in European samples in the UKB, stratified by sex and age groups (x-axis). Vertical black lines on top of the bars in panel (**B**) show analytical standard errors for the $R^2$ metric.

### S2 Supplementary Tables

| Phenotype | Phenotype Code | EFO Code | PGS | PGS Catalog ID | Training cohort | Notes |
|---|---|---|---|---|---|---|
| Standing Height | HEIGHT | EFO_0004339 | PGS002800 | PGS002800 | CSA | |
| | | | PGS002801 | PGS002801 | AFR | |
| | | | PGS002803 | PGS002803 | EAS | |
| | | | PGS002804 | PGS002804 | EUR | |
| | | | PGS002805 | PGS002805 | AMR | |
| LDL Cholesterol | LDL | EFO_0004611 | PGS000886 | PGS000886 | AFR | |
| | | | PGS000890 | PGS000890 | EAS | |
| | | | PGS000892 | PGS000892 | EUR | |
| | | | PGS000894 | PGS000894 | AMR | |
| | | | PGS000896 | PGS000896 | CSA | |
| HDL Cholesterol | HDL | EFO_0004612 | PGS003768 | PGS003768 | EUR | |
| | | | PGS003770 | PGS003770 | AFR | |
| | | | PGS003775 | PGS003775 | EAS | |
| | | | PGS003780 | PGS003780 | CSA | |
| Log triglycerides | LOG_TG | EFO_0004530 | PGS003802 | PGS003802 | EUR | |
| | | | PGS003804 | PGS003804 | AFR | |
| | | | PGS003809 | PGS003809 | EAS | |
| | | | PGS003814 | PGS003814 | CSA | |
| Testosterone | TST | EFO_0004908 | PGS_TST_M | | Male | Inferred using VIPRS. Sumstats: `https://zenodo.org/records/7222725` |
| | | | PGS_TST_F | | Female | |
| Urate | URT | EFO_0004531 | PGS_URT_M | | Male | Inferred using VIPRS. Sumstats: `https://zenodo.org/records/7222725` |
| | | | PGS_URT_F | | Female | |
| Creatinine | CRTN | EFO_0004518 | PGS_CRTN_M | | Male | Inferred using VIPRS. Sumstats: `https://zenodo.org/records/7222725` |
| | | | PGS_CRTN_F | | Female | |

Table S1: A listing of the quantitative phenotypes analyzed in this manuscript, along with the stratified polygenic scores used to predict them in the main analyses. Each phenotype is annotated with its Experimental Factor Ontology (EFO) code. The PGSs are identified with their PGS Catalog ID [3], where applicable, as well as the training cohort from which they were inferred.

| **Ancestry code** | **Definition** | **UKB Sample Size** | **CaG Sample Size** |
|---|---|---:|---:|
| AFR | African | 8,125 | 513 |
| AMR | Admixed American | 915 | 580 |
| CSA | Central and South Asian | 10,817 | 181 |
| EAS | East Asian | 2,704 | 259 |
| EUR | European | 460,416 | 26,813 |
| MID | Middle Eastern | 1,390 | 643 |
| OTH | Other | 2,189 | 344 |

Table S2: Sample sizes by predicted continental ancestry group in the UK Biobank (UKB) and CARTaGENE biobank (CaG) datasets. The ancestry prediction was based on a random forest classifier that takes as input top Principal Components of the genotype matrix and predicts continental ancestry, as defined in the 1000G+HGDP reference set [2]. These ancestry predictions are based on the `run_ancestry` procedure implemented in the `pgsc_calc` (v2.0.0) utility [4].

| **Prediction accuracy on Standing Height for AMR samples** | | |
|---|---|---|
| **UK Biobank** | Incremental $R^2$ | Pearson $R$ |
| CSA | 0.012 | 0.170 |
| AFR | 0.054 | 0.286 |
| EAS | 0.064 | **0.313** |
| EUR | **0.139** | 0.283 |
| AMR | 0.065 | 0.227 |
| **CARTaGENE** | Incremental $R^2$ | Pearson $R$ |
| CSA | 0.005 | 0.164 |
| AFR | 0.056 | **0.352** |
| EAS | 0.061 | 0.337 |
| EUR | **0.135** | 0.296 |
| AMR | 0.063 | 0.259 |

Table S3: Prediction accuracy of ancestry-stratified polygenic scores for Standing Height, evaluated on Admixed American (AMR) samples in two biobanks: UK Biobank and CARTaGENE. The names of the PGS models correspond to the continental ancestry of the samples from which it was inferred. The ancestry groups are Central and South Asian (CSA), African (AFR), East Asian (EAS), European (EUR), and Admixed American (AMR). The weights for the polygenic scores were obtained from the PGS Catalog (Publication ID: `PGP000382`). The accuracy metrics reported are incremental $R^2$ and Pearson correlation between the PGS and the phenotype.

| Ancestry | Total N | Prop. Male | Mean Age | HDL (mmol/L) | LDL (mmol/L) | TG (mmol/L) |
|---|---|---|---|---|---|---|
| EAS | 223,439 | 0.51 | 59.17 | 1.38 | 2.93 | 1.55 |
| EUR | 1,340,230 | 0.54 | 55.96 | 1.34 | 3.50 | 1.87 |
| AdmAFR | 101,272 | 0.66 | 53.99 | 1.16 | 3.37 | 1.99 |
| HIS | 48,253 | 0.62 | 51.78 | 1.11 | 3.48 | 2.44 |
| SAS | 38,479 | 0.73 | 52.81 | 1.05 | 3.11 | 1.99 |

Table S4: Characteristics of the Global Lipids Genetics Consortium (GLGC) cohorts used for GWAS and polygenic score construction. This summary data is aggregated from Supplementary Table 1 (**ST.1 Contributing cohorts**) of Graham et al. (2021) [1]. Mean blood lipid level units were converted from mg/dL to mmol/L by the standard conversion factors of 38.67 for HDL/LDL cholesterol and 88.57 for triglycerides. The ancestry groups represented are `EAS` (East Asian), `EUR` (European), `AdmAFR` (Admixed African), `HIS` (Hispanic), and `SAS` (South Asian).
